# Molecular insights into Profilin1-dependent regulation of cellular phosphatidylinositol-(4,5)-bisphosphate

**DOI:** 10.64898/2025.12.22.695975

**Authors:** Andrew Orenberg, Michael Chirumbolo, Ian Eder, Jia-Jun Liu, Silvia Liu, David Gau, Yubo Tang, Klemens Rottner, Jianhua Luo, Gerald V. Hammond, Partha Roy

**Affiliations:** Bioengineering, University of Pittsburgh; Cell Biology, University of Pittsburgh; Pathology, University of Pittsburgh; Institute of Zoology, Technical University of Braunschweig

**Keywords:** Profilin1, PIP_2_, phospholipase, hydrolysis, diacylglycerol, actin

## Abstract

Phosphatidylinositol (4,5)-bisphosphate (PIP_2_), the most abundant cellular poly-phosphoinositide (PPI) class of phospholipid, is a central plasma membrane (PM)-associated signaling hub that controls many cellular processes. In this study, we demonstrate that either deletion of the gene encoding actin-binding protein profilin1 (Pfn1) or disruption of Pfn1-actin interaction leads to downregulation of PM PIP_2_ content in cells. This is also phenocopied when F-actin is depolymerized implying that Pfn1-dependent PIP_2_ alteration is related to its actin-regulatory function. Phospholipase C (PLC) activity is critical for Pfn1-deficient cells to exhibit the PIP_2_-related phenotype. These findings, taken together with biochemical signatures of elevated PIP_2_ hydrolysis (higher baseline PM diacylglycerol-to PIP_2_ ratio and protein kinase C activity) exhibited by Pfn1-deficient cells, imply that PLC-mediated PIP_2_ hydrolysis plays a role in Pfn1-dependent regulation of PM PIP_2_. Furthermore, we unexpectedly found that Pfn1 loss leads to dramatic alterations in several other important forms of lipids, revealing a previously unrecognized role of Pfn1 as a broad regulator of cellular lipid environment that extends beyond PPI control. In conclusion, our study establishes Pfn1 as an important regulator of cellular lipid homeostasis.

**SUMMARY STATEMENT:** This study uncovers a mechanism of how functional loss of Profilin1, a key regulator of actin cytoskeleton, can trigger downregulation of plasma membrane content of PIP_2_, an important class of phospholipid, in cells.

## INTRODUCTION

Phosphatidylinositol (4,5)-bisphosphate (PI(4,5)P₂ - hereon referred to as PIP_2_) is a poly-phosphoinositide (PPI) class phospholipid that plays a central role in the regulation of diverse cellular processes. Although it constitutes a minor component of the inner leaflet of the plasma membrane (PM), PIP₂ is the most abundant PPI and serves as a critical signaling molecule and structural regulator in cells (reviewed in (Czech, 2000; Dickson and Hille, 2019; Mandal, 2020)). For example, its hydrolysis by phospholipases (PLCs) generates important second messengers, namely inositol trisphosphate (IP₃) and diacylglycerol (DAG), to promote the intracellular release of calcium and activation of protein kinase C (PKC), respectively. Metabolic conversion of PIP_2_ into D3-PPIs (e.g. PI(3,4,5)P_3_ [PIP_3_] and PI(3,4)P_2_) and negative regulation of those pathways involving the actions of various site-specific kinases (e.g. phosphatidylinositol 3’-kinase [PI3K]) and phosphatases (e.g. SHIP2 (SH2-domain containing inositol phosphatase 2, PTEN (phosphatase and tension homolog deleted on chromosome 10), and INPP4 (inositol polyphosphate-4-phosphatse) also influences pathways of cell proliferation, survival, and metabolism. Additionally, PIP_2_ directly interacts with a broad array of proteins involved in cytoskeletal remodeling (elaborated below), influencing a host of other processes including membrane trafficking (Thapa and Anderson, 2012), ion channel regulation (Harraz et al., 2020), and cell-cell/cell-extracellular matrix (ECM) adhesion (Akiyama et al., 2005; Chinthalapudi et al., 2014). Through these interactions, PIP₂ helps orchestrate dynamic cellular responses to environmental stimuli and maintains the spatial organization of signaling domains. Given its centrality in coordinating membrane-associated events, precise regulation of PIP₂ synthesis, localization, and turnover is essential for normal cellular function and viability.

Many important actin-binding proteins (ABPs) are influenced by their direct interactions with PIP_2_. For example, simultaneous binding of PIP_2_ and the small GTPase Cdc42 to the Wiskott-Aldrich Syndrome protein (WASP) and its homolog N-WASP induces conformational activation of these proteins (Higgs and Pollard, 2000). This activation promotes their interaction with the Arp2/3 complex, initiating actin polymerization. PIP_2_ has also been shown to sequester cofilin, an ABP responsible for actin filament disassembly, thereby inhibiting its depolymerizing activity and further enhancing actin filament stability (Zhao et al., 2010). Beyond direct modulation of actin filament dynamics, PIP_2_ also influences cytoskeleton-associated signaling pathways. Notably, it releases the autoinhibited conformation of talin, an ABP that links the actin cytoskeleton to the plasma membrane (PM), thereby facilitating integrin-mediated adhesion to the ECM (Ye et al., 2016). In a similar manner, PIP_2_ can facilitate the interaction with focal adhesion-associated protein vinculin with Arp2/3 complex to initiate actin nucleation at the sites of nascent adhesions (Braun et al., 2021; James et al., 2025). These findings underscore a central role of PIP_2_ in actin cytoskeletal regulation.

Profilins (Pfns) belong to a small family of G-actin-binding proteins. Pfn facilitates the nucleotide exchange from ADP- to-ATP-bound G-actin and delivers ATP-actin directly to the barbed ends of growing actin filaments. By virtue of their affinity for poly-L-proline (PLP) motifs, Pfn also interacts with and delivers G-actin to several important PLP-domain-bearing actin assembly-inducing factors (example: N-WASP, formins, Ena/VASP) to further enhance their ability to polymerize actin (Davey and Moens, 2020; Ding et al., 2012). These functions enable Pfn as a critical regulator of actin cytoskeletal dynamics in cells. Pfn also interacts with various PPIs (Lambrechts et al., 1997; Lassing and Lindberg, 1985) although the physiological significance of Pfn-PPI interaction has been largely understudied particularly in cellular settings. Lipid micelle- and/or model membrane-based studies have shown that Pfn1 (the most abundantly expressed cellular isoform of Pfn and the focus of the study) binds to PIP_2_ as well as PPIs that are generated downstream of PIP_2_ (such as PIP_3_ and PI(3,4)P_2_) (Lu et al., 1996; Sohn et al., 1995). Since PIP_2_ dissociates Pfn1 from G-actin, PIP_2_-binding has been postulated as a potential mechanism of negatively regulating sub-membranous Pfn1-actin interaction (Lassing and Lindberg, 1985); however, direct experimental evidence for this tenet in actual cells is still lacking in literature.

Complementing the widely accepted view of PIP_2_ acting as a regulator of ABP functionality in cells, two previous Pfn1-related studies from our group suggested that the reverse can be also true in a cellular setting. Specifically, we demonstrated that acute Pfn1 knockdown leads to plasma membrane (PM) enrichment of PI(3,4)P_2_ thereby influencing PI(3,4)P_2_-dependent PM recruitment of lamellipodin-VASP complex and cell migration (Bae et al., 2010), providing evidence for Pfn1-dependent modulation of cell behavior via PM PPI control. In a recent study we further showed that EGF-induced production of PIP_3_ (the PPI direct upstream of PI(3,4)P_2_) at the PM as well as the PM residence time of SHIP2 (the enzyme responsible for the metabolic conversion of PIP_3_ to PI(3,4)P_2_) in cells are enhanced in acute Pfn1 knockdown condition (Ricci et al., 2024). Although PIP_2_ serves as the immediate precursor PPI for PIP_3_ generation, our overexpression and knockdown experiments reported in that study collectively supported a premise that Pfn1 positively influences the basal cellular PIP_2_ content, and this is a generalizable feature across different cell types. The goal of the present study was to gain mechanistic insights into Pfn1-dependent regulation of cellular PIP_2_ content. This study provides the first direct evidence for PLC-mediated PIP_2_ hydrolysis as a mechanism for Pfn1-dependent regulation of PM PIP_2_ content in a cellular setting. However, Pfn1-dependent changes in PIP_2_ are driven by its role in actin dynamics rather than through direct PPI interaction. Our studies also unexpectedly identify Pfn1 as a broad regulator of cellular lipid environment that extends beyond PPI control.

## RESULTS

### Pfn1 depletion-induced PIP_2_ reduction is not an acute cellular response

In our previous study (Ricci et al., 2024), we showed that transient knockdown of Pfn1 expression in several different types of human cells (e.g. breast cancer cell lines: MDA-MB-231 (MDA-231) and BT-474; cervical cancer cell line: HeLa) led to an appreciable (35-50% depending on the cell type) reduction in PIP_2_ at the PM. Biochemical analyses of bulk lipids performed in MDA-231, BT-474, and MCF10A (a transformed mammary epithelial cell line) cells further indicated that total cellular PIP_2_ content is reduced upon transient Pfn1 loss. Conversely, we reported that overexpression of Pfn1 (either transiently or in a stable manner) leads to PIP_2_ enrichment. Given these findings, we first asked whether PM PIP_2_ reduction is maintained or somehow adaptively restored under complete and/or prolonged loss of Pfn1. To address this question, we performed immunofluorescence-based assessment of PM PIP_2_ in MDA-231 as well as B16F1 (a mouse melanoma cell line) cells that are engineered for CRISPR knockout (KO) of the *Pfn1* gene vs their respective control counterparts. Note that Pfn1 KO variant of B16F1 cells has permanent loss of Pfn1 expression serving us as a model system to study the impact of prolonged Pfn1 depletion on PIP_2_. In contrast, Cas9 expression in MDA-231 cells is doxycycline (dox)-inducible that allowed us to study the acute impact (5-6 days post-triggering of Cas9 induction) of complete loss of Pfn1 expression on PIP_2_. Immunoblot data shown in **Figure 1A** demonstrates the loss of Pfn1 expression in each of these two cell lines. As previously done in other studies (Hammond et al., 2006; Hammond et al., 2009), for PM PIP_2_ analysis, we specifically quantified the total rather than the average PM PIP_2_ staining intensity for three reasons. First, PIP_2_ is non-uniformly distributed across the PM, and therefore the average intensity calculation collapses a lot of biologically meaningful spatial information. Second, the average intensity calculation is impacted by significant cell shape and area differences that exist between cells within a group as well as between groups. Third, the integrated PM intensity is a better metric of how much total PIP_2_ is available for metabolic turnover on a cell-by-cell basis. Our PIP_2_ immunostaining experiments revealed that Pfn1 KO variants of both cell lines exhibited significantly lower total PM PIP_2_ content *vs* their respective control counterparts (**Figures 1B-C**). The average reductions in total PM PIP_2_ staining intensity in MDA-231 and B16F1 cells resulting from Pfn1 loss were equal to 45% and 20%, respectively (we offer potential explanations for quantitative differences in Pfn1-dependent PIP_2_ changes between the two cell lines in the discussion section). These data demonstrate that PM PIP_2_ reduction is maintained even under complete and/or prolonged loss of Pfn1 expression.

**Fig 1.**
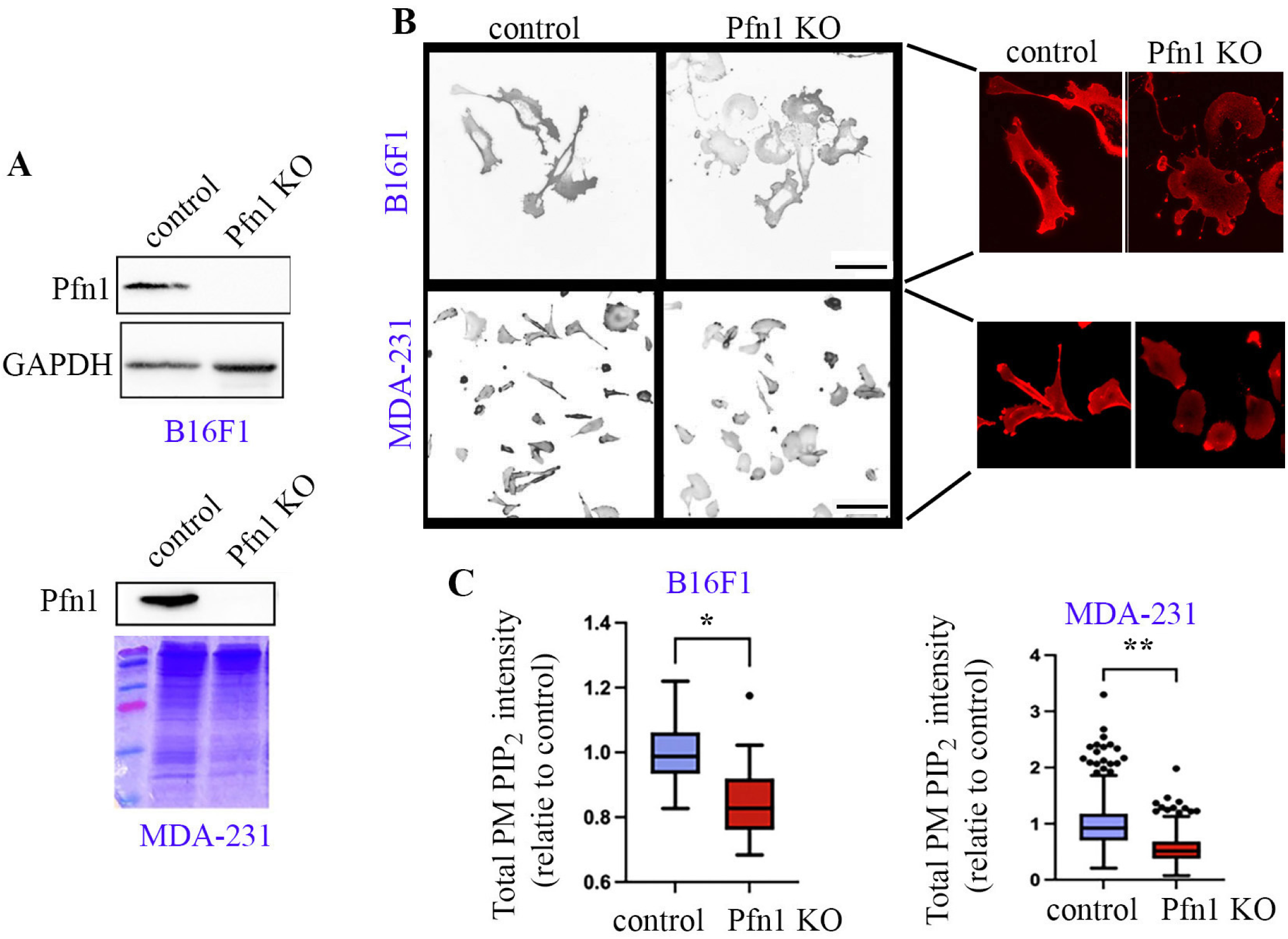
Pfn1 depletion-induced PIP_2_ reduction is not an acute cellular response. **A)** Representative Pfn1 immunoblots of cellular lysates from control and Pfn1 KO variants of B16F1 MDA 231 cells (GAPDH blot and Coomassie stained gels serve as loading controls) **B)** Representative confocal images of control and Pfn1 KO variants of B16F1 (60X objective) and MDA-231 (20X objective) cells immunostained for PIP_2_ (magnified images of selected regions are shown alongside). **D)** Box and Whisker plots summarizing the total PM PIP_2_ staining of Pfn1 KO cells relative to their control counterparts for each cell line. Each box represents the data falling within the 25^th^ to 75^th^ percentile range, and a line denoting the mean value of the data set. Whiskers represent 1.5x the interquartile range above and below the 25 and 75 percentiles. The data sets represent analyses of 200-300 and ∼400 cells pooled from three independent experiments for B16F1 and MDA-231 cell lines, respectively (*: p<0.05; **p: <0.001). Scale bars in the upper and lower sub-panels in panel B represent 100 μm and 40 μm, respectively.

### Disruption of actin-binding of Pfn1 phenocopies PIP_2_ phenotype of Pfn1-deficient cells

Next, to determine which functionality of Pfn1 is linked to its regulation of cellular PIP_2_, we performed functional rescue experiments where Pfn1 KO B16F1 cells were transfected with plasmids encoding myc-tagged versions of either wild-type (WT) Pfn1 or various ligand binding-deficient point-mutants of Pfn1 before performing PIP_2_ immunostaining. These mutants include: H119E-Pfn1 (∼50 fold deficient in actin-binding), H133S-Pfn1 (deficient in PLP-binding), and R88L-Pfn1 (deficient in PPI-binding). Note that due to structural overlap between the actin- and PPI-binding regions of Pfn1, R88L-Pfn1 has a 2-fold reduction in actin-binding (Bae et al., 2010). While PIP_2_-binding of WT vs H119E-Pfn1 has never been directly quantified in biochemical assays, we previously showed that H119E substation does not affect the membrane fraction of ectopically overexpressed Pfn1 in cells (Bae et al., 2010). Furthermore, in purified protein-lipid vesicle mixture settings, an analogous mutant H119D-Pfn1 inhibits PLCγ-mediated PIP_2_ hydrolysis as efficiently as WT-Pfn1 (Sohn et al., 1995). These findings suggest that H119D/E-Pfn1 retains intact membrane PPI binding. All four rescue Pfn1 constructs were expressed in an IRES (internal ribosomal entry site)-GFP backbone vector that allowed identification of rescued cells by the presence of GFP fluorescence. As additional transfection controls, both control and Pfn1 KO cells were transfected with an IRES-GFP backbone vector. Surprisingly, PIP_2_ immunostaining revealed that PM PIP_2_ reduction in Pfn1-deficient cells is reversible by all Pfn1 rescue constructs except the actin-binding deficient mutant H119E-Pfn1 (**Figures 2A-B**). Since GFP and Pfn1 rescue constructs are linked by an IRES, we analyzed GFP fluorescence intensity of cells selected for PIP_2_ analyses as a surrogate measure for comparing the relative expressions of Pfn1 rescue constructs across the various groups. As per these analyses, the average GFP expression of cells chosen for PIP_2_ analyses was found to be comparable between the various Pfn1 KO-rescue groups (**Figure 2C**). Therefore, we argue that our observed phenotypic differences related to PIP_2_ are not confounded by the expressions of various Pfn1 rescue constructs. Based on these results, we conclude that Pfn1-dependent changes in PIP_2_ are primarily attributed to its actin-related function.

**Fig 2.**
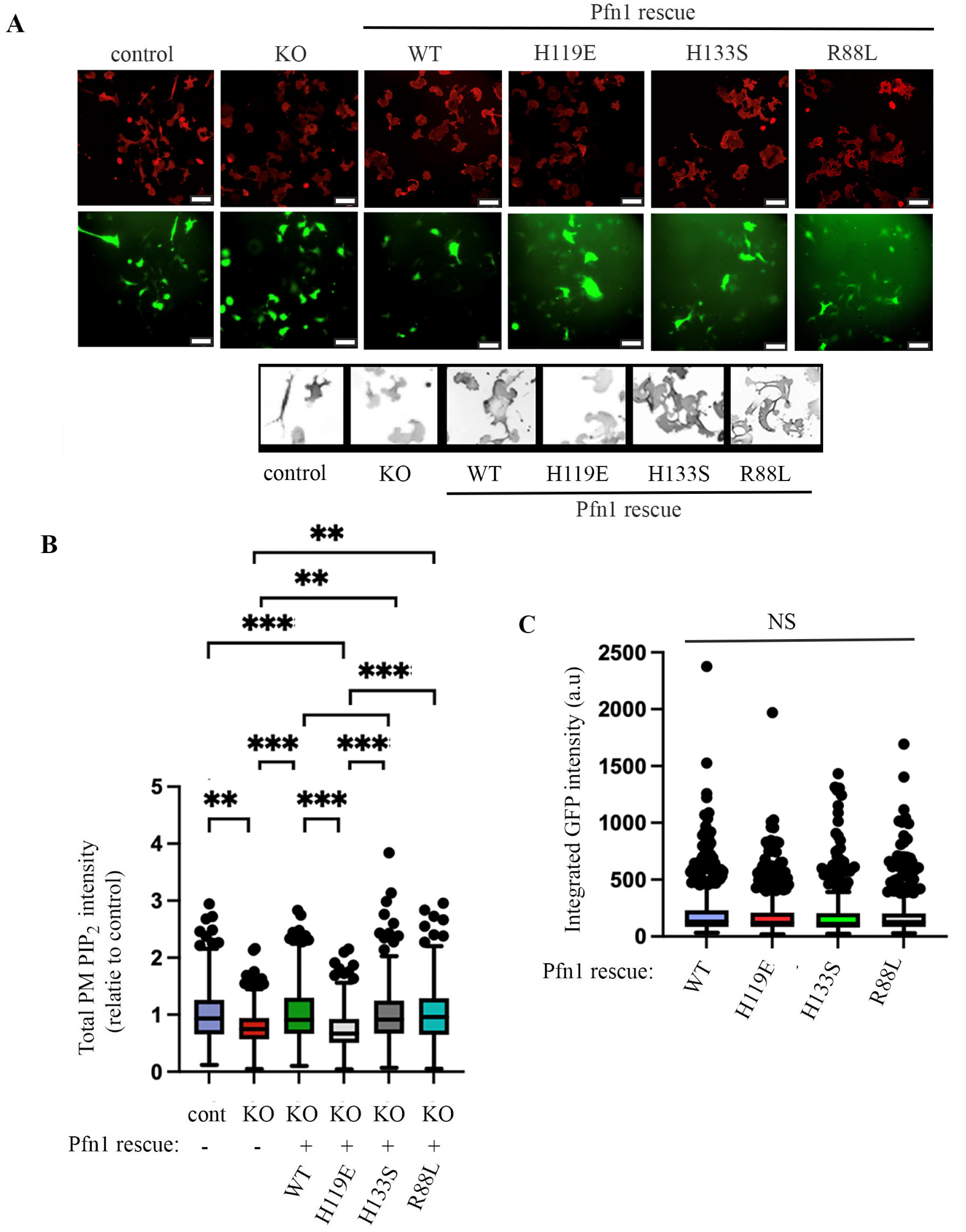
Disruption of actin-binding of Pfn1 phenocopies the PIP_2_ phenotype of Pfn1-deficient cells. **A)** Representative 20X imaging of PIP_2_ immunostaining (red) of control and Pfn1 KO B16F1 cells with or without functional rescue by wild-type (WT) and the indicated mutants of Pfn1 GFP-positive cells indicate the transfected cells; empty vector GFP transfection serves as the control). (magnified PIP_2_ staining images of selected GFP+ cells in each group are shown in the bottom). Scale bar represents 100 μm. **B)** Quantification of PM PIP_2_ intensity in GFP-positive cells for various groups relative to control (control cells transfected with GFP). Each datapoint represents an independent experiment; a total of ∼300-450 cells were quantified pooled from three independent experiments (**: p<0.01; ***: p<0.001). **C)** Quantification of relative GFP expression for the various rescue groups (each data point is the mean readout of all GFP+ cells for a given experiment; NS – not significant).

Since Pfn1 loss generally leads to reduced level of polymerized actin in cells including B16F1 cells (**Figures 3A-B**), we next asked whether acute actin depolymerization could invoke a similar PIP_2_ loss in cells. To test this possibility, we performed PIP_2_ immunostaining of B16F1 cells subjected to acute treatment (5 min) of actin depolymerizing drug Latrunculin-B (LatB) vs DMSO (vehicle control). Cells were also counterstained with phalloidin to assess the changes in F-actin upon LatB treatment. Our experiments showed that global actin depolymerization by LatB treatment was also accompanied by a significant reduction in PM PIP_2_ content of B16F1 cells **(Figures 3C-D**). To further determine if there is any temporal correlation between LatB-induced F-actin disruption and PIP_2_ attenuation, we performed orthogonal live-cell imaging of HEK-293 cells co-transfected with PIP_2_ biosensor GFP-PH-PLCδ along with a PM marker (iRFP-Lyn11) and FastActX (a probe for F-actin) before and after acute treatment of LatB (DMSO treatment served as control). Consistent with our observations in B16F1 cells, LatB treatment resulted in a rapid decrease in the PM-to-cytosolic fluorescence ratio (indicative of PM PIP_2_ reduction) in HEK-293 cells that remarkably synchronized with actin depolymerization marked by the loss of F-actin fluorescence intensity (**Figures 3E-G**). On a kinetic scale, LatB-induced PIP_2_ decline occurred slightly slower and persisted longer than the F-actin response, with the mean half-lives of PIP_2_ and F-actin changes equal to approximately 2.2 min and 1 min, respectively (**Figures 3H**). In a separate experiment, we studied the impact of short-term (30 min) treatment of non-muscle myosin II inhibitor blebbistatin (vs DMSO as vehicle control) on PM PIP_2_ content in MDA-231 cells (**supplementary Fig S1**). The major effects of blebbistatin on actin cytoskeleton are disintegration of actin stress fibers, softening of cortical actin, and transformation of lamellipodial actin into loose network of accumulated amorphous actin structures that correspond to membrane ruffles (Shutova et al., 2012). These phenotypes were also recapitulated in our experimental settings (**supplementary Figure S1A**). In general, blebbistatin-treated cells exhibited protrusive structures in random directions with PIP_2_ enrichment in peripheral F-actin-rich regions (consistent with the LatB experimental data) and a higher (p=0.09) overall cell edge PIP_2_ staining vs vehicle-treated cells, further underscoring a prominent impact of actin cytoskeletal perturbation on PM PIP_2_ (**supplementary Figures S1B-C**). Collectively, the findings from our Pfn1 KO-rescue and various actin-related pharmacological studies suggest that Pfn1-dependent PIP_2_ alteration is likely a secondary consequence of peripheral actin cytoskeletal disturbances in cells.

**Fig 3.**
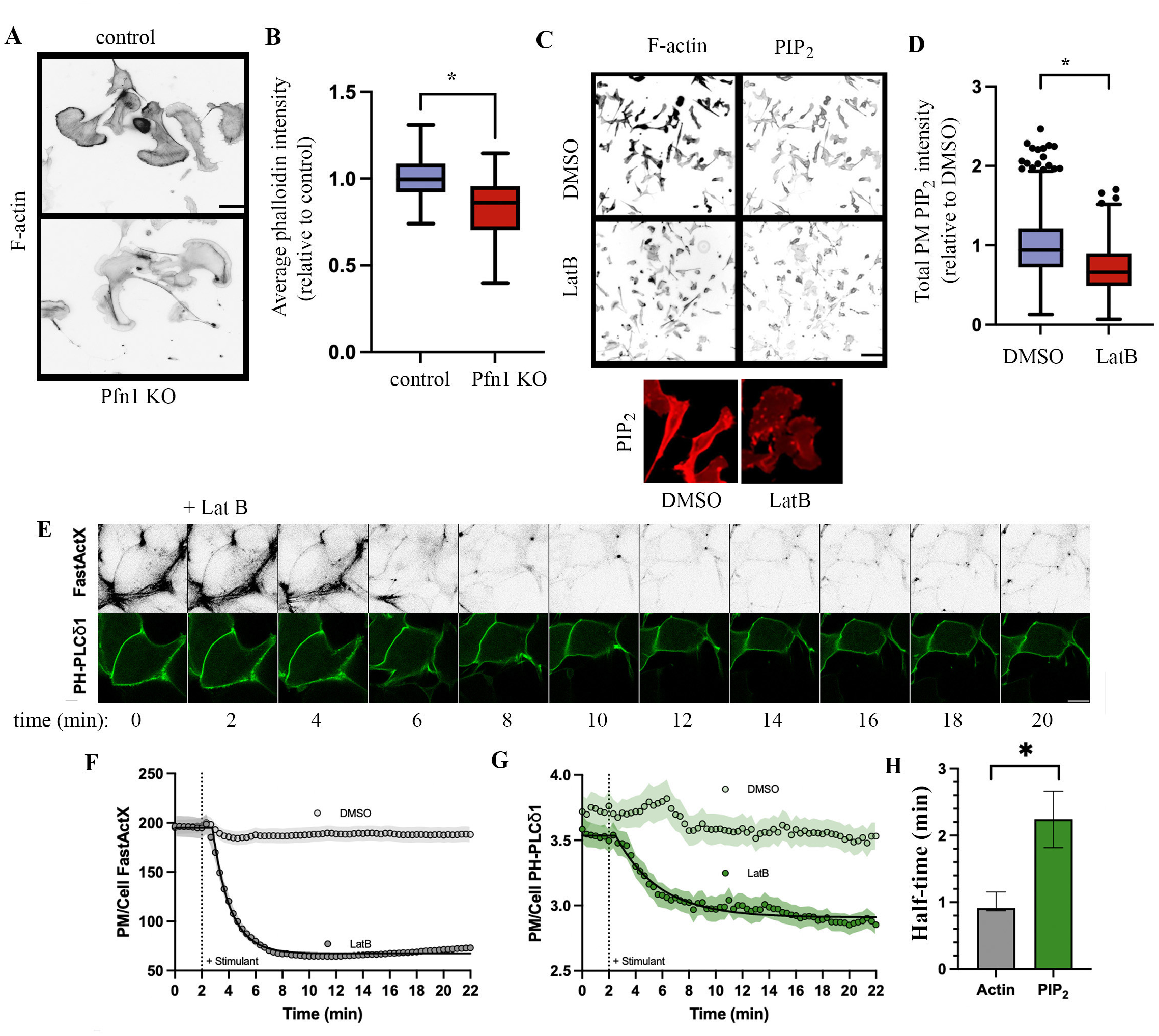
Actin depolymerization triggers rapid loss of PM PIP_2_. **A-B)** Representative 60X fluorescence images (panel A) of control and or Pfn1 KO B16F1 cells stained with phalloidin. Panel B shows the associated quantification with each dataset representing quantification of ∼45 fields pooled from three independent experiments. Scale bar represents 20 μm. **C-D**) Representative 20X fluorescence images (panel C) of B16F1 cells stained with phalloidin and anti-PIP_2_ antibodies in DMSO (control) vs LatB-treated conditions (magnified images of PIP_2_ staining of selected regions of interest are shown below in panel C). The box and whisker plot in panel D summarizes the LatB-induced changes in PM PIP_2_ intensity (∼500 cells were quantified per treatment group pooled from 3 independent experiments; *: p<0.05). **E-H)** Representative confocal fluorescence time-lapse images of HEK-293 cells expressing GFP-PH-PLCδ and treated with FastActX before and after LatB treatment (panel E). Panels F and G show the summary of temporal changes of F-actin (by FastActX fluorescence) and PM PIP_2_ (by PM/cytoplasmic fluorescence of PH-PLCδ) in response to LatB vs DMSO as control (traces represent fluorescence mean + SEM of all cells (DMSO – 50 cells, LatB – 45 cells) pooled from 3 individual experiments. Panel H shows the relative half-times of the F-actin and PIP_2_ response curves (*: p<0.05).

### PIP_2_ attenuation upon loss of Pfn1 is reversible by PLC inhibition

Based on our observation that PM PIP_2_ loss ensues rapidly (within minutes) in response to triggering actin depolymerization, we speculated that Pfn1-dependent changes are PIP_2_ could be somehow related to perturbation of its turnover rather than its synthesis pathway. PIP_2_ is turned over by two major pathways. One of these pathways involves PI3K-mediated conversion of PIP_2_ to PIP_3_ (followed by sequential conversion of PIP_3_ to PI(3,4)P_2_). Our previous studies showed evidence for enhanced EGF-induced production of D3-PPIs (i.e. PIP_3_ and PI(3,4)P_2_) in MDA-231 and HeLa cells upon Pfn1 depletion (Bae et al., 2010; Ricci et al., 2024). However, these D3-PPIs are transiently generated in response to acute activation of receptor tyrosine kinases (RTKs - EGFR, PDGFR, IGFR) and only a small fraction of the PM pool of PIP_2_ is converted into PIP_3_. Therefore, we reasoned that increased conversion of PIP_2_ to PIP_3_ is unlikely to explain ∼40% decrease in the basal PIP_2_ level observed in those cell lines upon Pfn1 depletion in steady-state serum-containing culture. A second pathway of PIP_2_ turnover involves hydrolysis of PIP_2_ into DAG and IP_3_ by the action of PLC class of enzymes. Therefore, to test whether pharmacological PLC inhibition has any effect on Pfn1-dependent changes in the PM PIP_2_ content, we performed PIP_2_ staining of control and Pfn1 KO MDA-231 cells following overnight treatment with either U73122 (a broad-spectrum PLC inhibitor) or its inactive analog U73343 (a negative control). As expected, in inactive compound (U73343) treatment setting, Pfn1 KO cells exhibited lower PM PIP_2_ content vs their control counterparts. However, when treated with the active PLC inhibitor U73122, the total PM PIP_2_ content was found to be conspicuously elevated in Pfn1 KO cells abolishing the PIP_2_ differential between the two groups of cells (**Figures 4A-B**). The two major classes of PLC enzymes that hydrolyze PIP_2_ are PLCγ (activated downstream of RTKs) and PLCβ (activated downstream of G protein–coupled receptors (GPCRs)). In model membrane studies utilizing purified proteins (Pfn1, PLCγ), Goldschmidt-Clermont and colleagues previously demonstrated that Pfn1 inhibits PLCγ-mediated hydrolysis of PIP_2_ but this inhibitory action of Pfn1 on PIP_2_ hydrolysis is abolished when PLCγ is tyrosine-phosphorylated (mimicking the activated state of PLCγ in response to RTK activation) (Goldschmidt-Clermont et al., 1991; Goldschmidt-Clermont et al., 1990). In fact, in our previous study, we also failed to see any effect of Pfn1 depletion on the production of IP1 (an immediate metabolic byproduct of IP_3_) in MDA-231 cells in an acute EGF-stimulation setting (Ricci et al., 2024) suggesting that Pfn1’s ability to impact PIP_2_ hydrolysis by RTK-activated PLCγ is unlikely. Since PLCβ activity is also stimulated by active ingredients of serum (e.g. various bioactive lipids, hormones, and peptides), we next examined the effect of silencing the expression of PLCβ3 (the dominant isoform of PLCβ in MDA-231 cells) on PIP_2_ content in control vs Pfn1-deficient MDA-231 cells. Consistent with our pharmacological inhibition data, these experiments also revealed that silencing PLCβ3 expression increased PIP_2_ selectively in Pfn1-deficient cells, mitigating the PIP_2_ differential between the two cell types (**Figures 4C-E**). Collectively, these findings demonstrate that intact PLC activity is critical for Pfn1-deficient cells to exhibit the PIP_2_-related phenotype.

**Fig 4.**
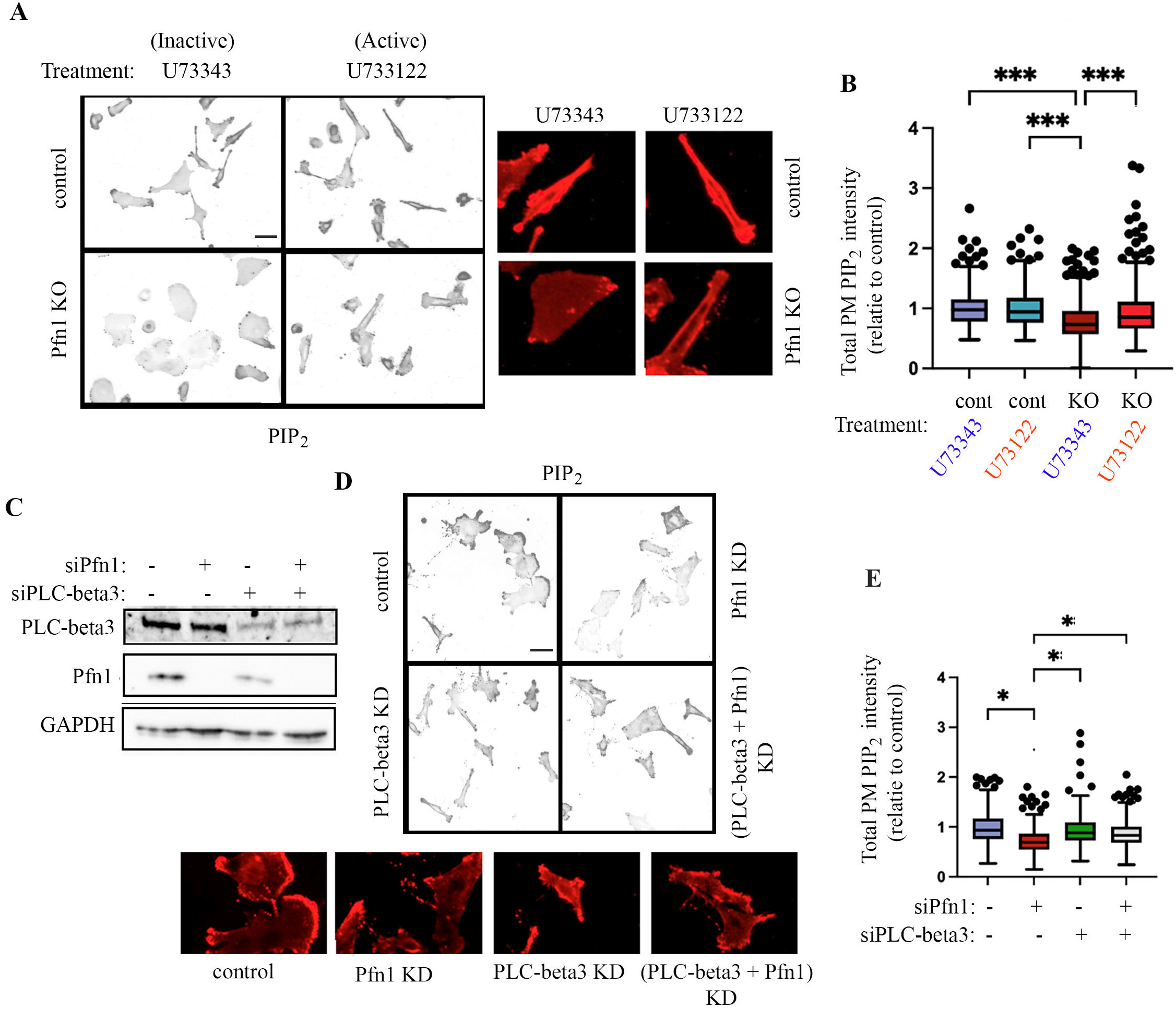
PIP_2_ attenuation upon loss of Pfn1 is reversible by PLC inhibition. **A-B)** Representative PIP_2_ staining images (*panel A;* 40X) of control and Pfn1 KO MDA 231 cells, treated with either inactive (U73343) or active (U73122) pan-PLC inhibitor drug (magnified images of PIP_2_ staining of selected regions of interest are shown alongside). Panel B shows the associated PM PIP2 quantification for the various groups (all data normalized to the average readout of control cells treated with the U73343 compound. Each datapoint represents an independent experiment; a total of ∼ 450 cells/group were analyzed from three independent experiments. Scale bar represents 40 μm**. C-E)** *Panel C* - Representative PLCβ3 and Pfn1 immunoblots of lysates of MDA-231 cells following transient transfection of the indicated siRNAs (non-targeting control siRNA transfection served as control). C) GAPDH blot serves as the loading control. Representative PIP_2_ staining images fluorescence images (*panel D;* 40X; scale bar – 80 μm) of MDA-231 cells transfected with the indicated siRNAs (magnified images of PIP_2_ staining of selected regions of interest of each group are shown underneath), and the associated quantification (*panel E*) to compare the PM PIP_2_ intensity between the various groups of cells based on the results of three independent experiments (*: p<0.05).

### Pfn1-deficient cells exhibit signatures of elevated PIP_2_ hydrolysis

The results of our PLC inhibition studies can be interpreted in at least two different ways. One possible scenario is that Pfn1’s presence somehow acts as a brake on PLC-mediated PIP_2_ hydrolysis, and this inhibition is lost in Pfn1-deficient cells resulting in PM PIP_2_ loss through accelerated PIP_2_ hydrolysis. Alternatively (but not necessarily in a mutually exclusively manner), the signaling downstream of activated PLC is somehow indirectly responsible for Pfn1-dependent modulation of PIP_2_. We asked whether Pfn1-depleted cells bear any of the molecular signatures of increased PIP_2_ hydrolysis. To address this, we first performed quantitative mass-spectrometry (MS)-based lipidomic analyses of control vs Pfn1 KO MDA-231 cells which revealed several interesting findings. First, the total lipid content (normalized to the total protein content) of Pfn1 KO cells was found to be ∼10% lower than control cells, and this difference was found to be significantly different (**Figure 5A**). Second, analyses of differentially abundant lipids (out of a total of ∼900 lipids) revealed that 89 and 104 lipids were upregulated and downregulated, respectively, in Pfn1 KO cells relative to their control counterparts (**Figure 5B**), suggesting that Pfn1 loss has a global impact on lipid profile that extends beyond PPI control. Differentially abundant lipids categorized in major classes are shown as a heat-plot in **Figure 5C**. Major lipids that were altered upon loss of Pfn1 expression in a statistically significant manner include phosphatidylcholine (PC; reduced by ∼25%), cholesterol (reduced by ∼17%), free fatty acid (FFA; reduced by ∼26%), sphingomyelin (reduced by ∼23%), and triglycerides (TGs: increased by ∼12%) (**Figures 5D-E).** Although the lipidomic panel utilized in our studies did not probe for PPIs, the PI content was found to be comparable between control and Pfn1 KO cells. Third, in 4 out of 5 experiments, we saw a general trend of elevated DAG (a direct hydrolysis product of PIP_2_) content in Pfn1 KO samples; however, the large experiment-to-experiment variability in the absolute content as well as Pfn1-dependent changes in DAG precluded us from achieving statistical significance between the two groups. The large variability in the measured DAG content in our experiments is not totally surprising since cellular DAG level is known to fluctuate with growth and/or impacted by unintended changes in the chemical parameters of culture conditions (Mondal et al., 2024; Van Veldhoven and Bell, 1988). Phosphatidic acid (PA) is a lipid byproduct of DAG that is generated by direct conversion of DAG by DGK (DAG-kinase) to replenish the pool of cellular PIP_2_ through a feedback signal. Encouragingly, we observed a near-statistically significant (p=0.06) increase in the absolute content of PA in Pfn1 KO cells relative to their control counterparts (**Figures 5D-E**). Note that although PA can be also generated by phospholipase-D (PLD)-mediated conversion of PC (a lipid that is also decreased dramatically in Pfn1 KO cells), it is highly unlikely that PA increase in Pfn1-deficient cells is reflective of increased PLD-mediated conversion of PC for two reasons. First, the observed magnitude of change in the content of PA (∼3000 pmol/mg increase) was disproportionately lower than that of PC (>200,000 pmol/mg decrease) in response to Pfn1 KO. Second, monomeric actin directly binds to and inhibits the activity of PLD (Lee et al., 2001). Therefore, increased G-to-F-actin ratio in Pfn1-deficient cells, if at all, would be expected to result in diminished PLD activity and PLD-mediated conversion of PC to PA.

**Fig 5:**
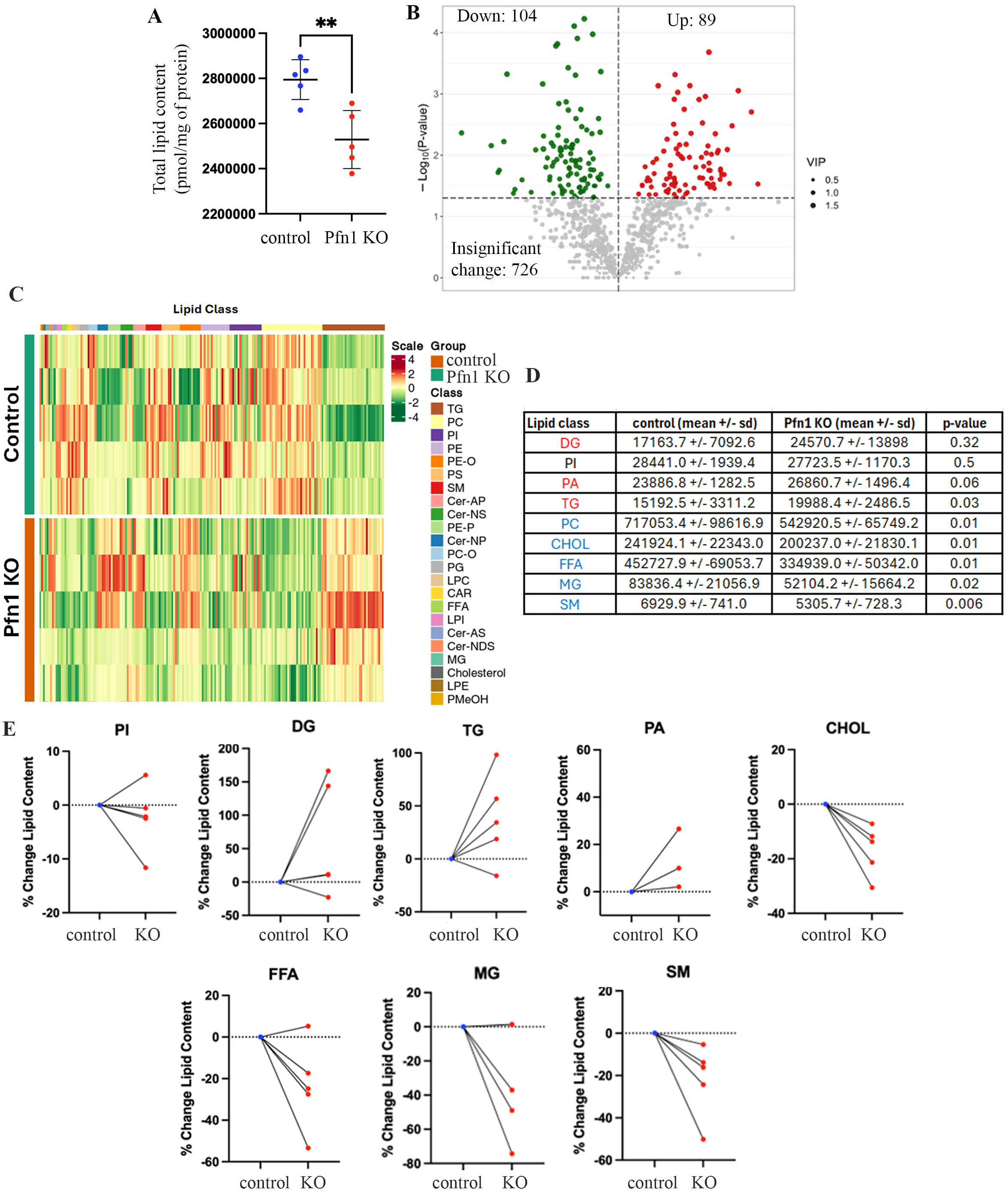
Pfn1 loss has a broad impact on cellular lipids. **A)** Quantification of the total lipid content (in pmol/mg of total protein) based on mass-spectrometry based on lipidomic analyses of control and Pfn1 KO MDA-231 cells (5 biological replicates/group with >1,000,000 cells per sample were analyzed). **B)** Volcano plot represents the number of differentially abundant lipid moieties measured in the lipidomic panel. **C)** Heat-plot showing directional alterations in various lipid classes upon loss of Pfn1 in MDA-231 cells. **D)** Quantitative comparison of the abundance (in pmlo/mg of total protein) of the indicated lipids between control and Pfn1 KO cells (PC - phosphatidylcholine, CHOL - cholesterol, FFA - free fatty acid, SM - sphingomyelin, TG - triglyceride; PI = phosphatidylinositol, PA - phosphatidic acid, and DAG – diacylglycerol; lipids in red and blue exhibited trends in up- and down-regulation, respectively, upon Pfn1 KO). **E**) Line plots showing % changes in the indicated lipids in Pfn1 KO relative to control cultures in all five biological replicates (note: near identical % changes in certain biological replicates led to overlapping line plots for some lipids).

While the general trends of elevated DAG and PA in Pfn1 KO cells are conceptually consistent with a scenario of increased PIP_2_ hydrolysis, a key limitation of whole-cell lipidomic analysis is that it fails to distinguish lipid changes in specific subcellular locations/compartments. This is particularly a relevant issue for DAG since: 1) the largest pool of DAG is in the endoplasmic reticulum (ER)/Golgi, 2) DAG is under constant metabolic flux, and 3) importantly, subtle changes in DAG resulting from PM PIP_2_ hydrolysis may be difficult to capture by whole cell lipidomic measurements. To circumvent these issues, we performed a series of additional studies to gain further biological insights into Pfn1’s effect on PIP_2_ hydrolysis as described in the following sections.

First, a key downstream signature of PLC-mediated PIP_2_ hydrolysis is DAG-mediated activation of serine-threonine kinase protein kinase C (PKC). Therefore, to assess Pfn1-dependent changes in PKC activation, we performed immunoblot analyses of total cell extracts prepared from control vs Pfn1-deficient MDA-231 cells with a phospho-PKC (pPKC) substrate antibody that recognizes all PKC-phosphorylated proteins. For these studies, cells were either serum-starved (representing unstimulated “baseline” condition) or subjected to acute stimulation of lysophosphatidic acid (LPA), a bioactive lipid and an active ingredient of serum that stimulates PLCβ activity via GPCR-activation. The baseline (i.e. in an unstimulated condition) PKC activity signature (measured by the intensity of pPKC-substrate immunoreactive bands) was clearly more pronounced in Pfn1-depleted MDA-231 vs their control counterparts (**Figures 6A-B**), a finding that was also reproducible in B16F1 cells (**Figure 6C-D).** When we compared LPA-induced increase in the PKC activity readout relative to the baseline condition, the fold-change values (control cells: ∼3 fold increase from baseline; Pfn1-deficient cells: ∼1.3-fold increase from baseline) implied that Pfn1 loss did not necessarily augment LPA-induced PKC activation response. A similar trend was observed when we analyzed LPA-induced Ca^2+^ response (another event downstream of PLC-mediated PIP_2_ hydrolysis triggered by IP_3_) by live-cell fluorescence imaging of GCaMP (a Ca^2+^ biosensor) transfected in MDA-231 cells (data summarized in **Figures 6E-I**). Both groups of cells exhibited rapid LPA-induced Ca^2+^ spike (reached its peak at ∼30 sec after stimulation) followed by a rapid decline reaching a post-stimulation resting level. To account for cell-to-cell variation in the actual expression of the biosensor, we baseline-corrected GCaMP fluorescence by normalizing each kinetic datapoint readout to the average pre-stimulation value on a cell-by-cell basis, for calculating the peak amplitude, integrated Ca^2+^ signal (area under the curve), and the post-stimulation resting value for each of the two groups. As per these analyses, we did not see any significant difference in either the peak amplitude or integrated Ca^2+^ signal between the control and Pfn1 knockdown groups, further underscoring the fact that Pfn1 loss does not necessarily confer cells an increased ability to respond to agonists (i.e. LPA-induced GPCR activation in this specific case). However, we noted that the post-stimulation resting Ca^2+^ signal was elevated in Pfn1-deficient cells relative to control cells (p<0.01), a feature that could result from increased basal PIP_2_ hydrolysis and/or reduced re-uptake of cytosolic Ca^2+^ by endoplasmic reticulum and/or reduced efficiency of Ca^2+^ export.

**Fig 6:**
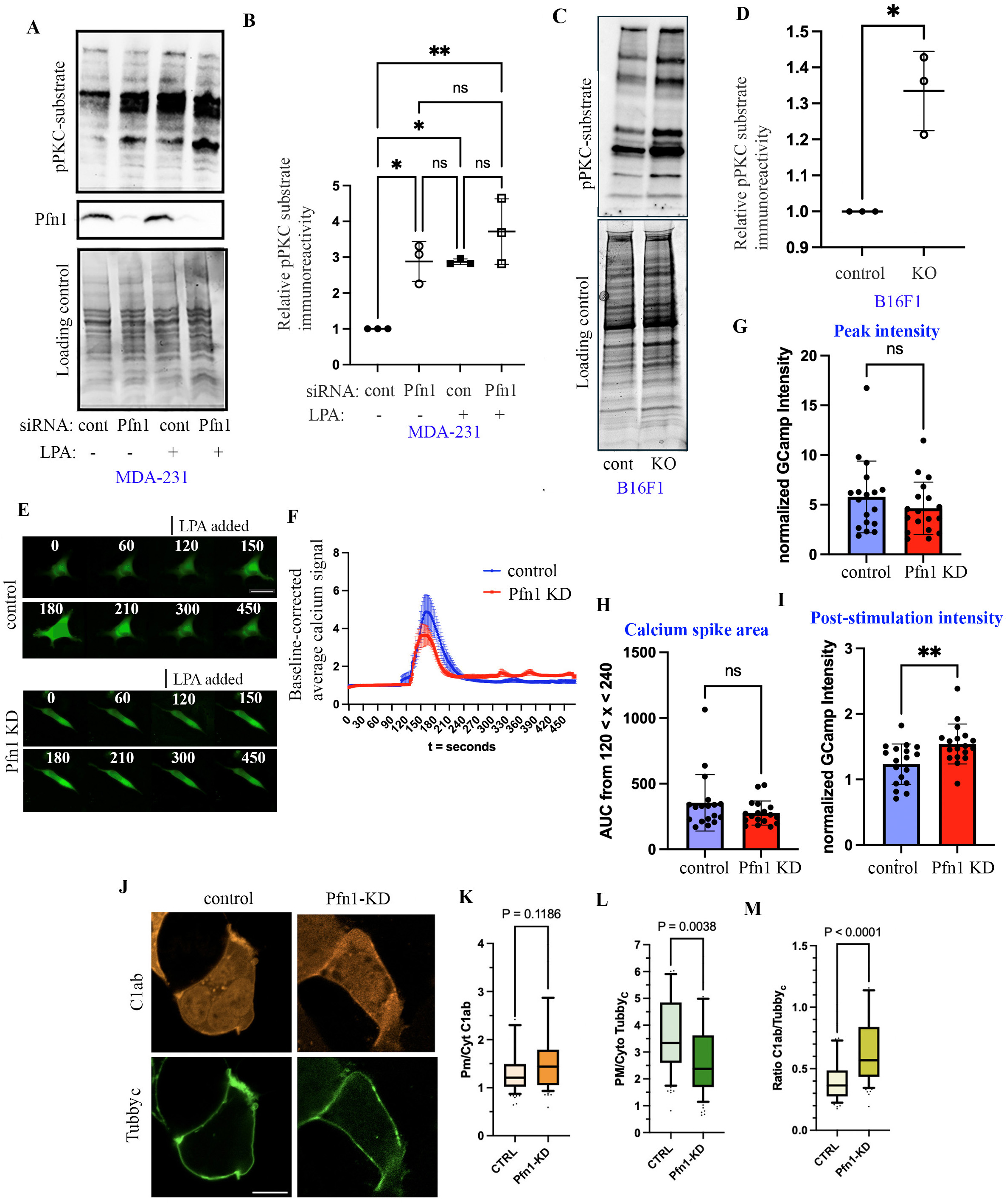
Supportive evidence for Pfn1-dependent changes in PIP_2_ hydrolysis signature. **A-B)** Representative phospho-PKC (p-PKC) substrate and Pfn1 immunoblots of extracts prepared from control and Pfn1-depleted MDA-231 cells (*panel A*) following treatment with either 10μM LPA or vehicle control (unstimulated) for 5 minutes; the stain-free gel image serves as the loading control (quantification of immunoblot results are shown in panel B; n=3 experiments). **C-D**) Representative baseline p-PKC substrate immunoblot of control vs Pfn1 KO B16F1 cultures (*panel C*) and the associated quantification (panel D; n=3 experiments). **E-I**) Representative live fluorescence images (*panel E; scale –* 20 μm) of MDA 231 cells with or without siRNA mediated depletion of Pfn1 (*panel E*), transfected with calcium ion probe GCaMP, treated with 10µM of LPA over 8 minutes with 2 minutes of baseline imaging and 6 minutes of imaging post treatment (stimulation at t = 120 seconds). Images depict t = 0, 60, 120, 150, 180, 210, 300, and 450 seconds. Panel F depicts the quantification of average baseline-corrected GCaMP integrated fluorescence intensity over time normalized to the average of 2 minutes pre-stimulation of the treatment groups. Panels G through I show the comparisons of peak Ca2+ indicator intensity, calcium spike area, and the post-stimulation intensity (relative to the pre-stimulation value) between the two groups. The dataset represents analyses of 18 cells/group pooled from 3 independent experiments; ns – not significant; **: p<0.01). **J-M**) Lipid biosensor studies demonstrating elevated DAG:PIP_2_ levels in HEK-293 cells in response to Pfn1 knockdown (KD) - Panel J shows representative confocal images of HEK-293 cells expressing Tubby_c_-EGFP PIP_2_ and mCherry-C1ab-PKD1 DAG biosensors, after treatment with non-targeting (ctrl) or Pfn1-directed siRNA pools (scale bar = 10 µm). Panels K through M show quantification of the PM to cytosolic intensity of the DAG and PI(4,5)P_2_ biosensors, as well as the ratio of the two measurements. Line, box and whiskers show the median, 25^th^, 75^th^, 10^th^ and 90^th^ percentiles of 76 (ctrl) or 74 (Pfn1 KD) cells from three independent experiments (P values show the results of Welch’s t-test).

Second, we performed lipid biosensor studies in HEK-293 cells to assess the impact of Pfn1 knockdown on the relative DAG-to-PIP_2_ content at the PM as a measure of intrinsic PIP_2_ hydrolysis efficiency (data summarized in **Figures 6J-M**). For these studies, HEK-293 cells were co-transfected with GFP-PH-Tubby_C_ (a C-terminal fragment of Tubby protein that binds to PM PIP_2_ and therefore, acts as a PIP_2_ biosensor) and mCherry-C1ab (a construct with tandem DAG-binding C1A-C1B domains of PKD1 that serves as a DAG biosensor) along with a PM marker (iRFP-Lyn11). We apriori confirmed that Pfn1 depletion by siRNA transfection did not significantly change the PM PI4P (the direct precursor PPI for PIP_2_ biosynthesis) content in HEK-293 cells (**supplementary Figure S2**), mirroring our previous findings in HeLa cells (Ricci et al., 2024). However, it resulted in reduced PM PIP_2_ content as indicated by a lower PM-to-cytoplasmic fluorescence ratio of GFP-Tubby_C_ in Pfn1 knockdown vs control cells (**Figure 6L**). The absolute PM-to-cytosolic fluorescence ratio of the DAG reporter was slightly higher in Pfn1 knockdown cells than control cells, but this was not statistically significant (**Figure 6K**). However, the difference was more pronounced and statistically significant when these DAG biosensor readouts were normalized to the PM-to-cytosolic ratio of PIP_2_ biosensor readouts on a cell-by-cell basis before comparison (**Figure 6L**). These data indicated higher DAG-to-PIP_2_ ratio (a measure of intrinsic PIP_2_ hydrolysis efficiency) at the PM in Pfn1 knockdown cells, Third, we performed gene set enrichment analyses of bulk transcriptomic data (summarized in **supplementary Figure S3**) of control vs Pfn1 KO MDA-231 cells. These analyses predicted enrichment of cytosolic IP_3_/IP_4_ (downstream metabolites of PIP_2_ hydrolysis) synthesis-related genes along with transcriptional upregulation of two PLC enzymes (PLCβ1 and PLCβ4) by ∼4-fold in Pfn1 KO cells. Collectively, these findings support Pfn1’s negative impact on PIP_2_ hydrolysis at the PM.

## DISCUSSION

In our previous study, we first reported a phenomenon that perturbing Pfn1 expression has a profound consequence on the cellular PIP_2_ content; however, the underlying mechanism remained unclear. The present study reports several new findings that shed novel biological insights into how Pfn1’s presence may positively impact the PM PIP_2_ content. First, we reveal that complete loss of Pfn1, either triggered acutely (as in MDA-231 cells) or in a prolonged manner (as in B16F1 cells) also leads to reduced PIP_2_ level at the PM, recapitulating our previous observations in transient knockdown settings in various cell types. This suggests that Pfn1-dependent change in PIP_2_ is not a transient cellular response that can be compensated by other mechanisms in the long term. Second, based on findings from *in vitro* studies utilizing purified proteins (Pfn1, PLC) and PPI-containing model membrane (Goldschmidt-Clermont et al., 1991; Goldschmidt-Clermont et al., 1990), it has been previously postulated in the literature that Pfn1 can protect PIP_2_ from PLC-mediated hydrolysis through its direct binding to PIP_2_. However, this postulate was never directly tested in a cellular setting. Notably, our study unexpectedly reveals that Pfn1-dependent changes in PM PIP_2_ are linked to its actin-regulatory function, highlighting that *in vitro* systems do not adequately recapitulate the complex cellular context in which Pfn1 operates. Third, we demonstrate that Pfn1-deficient cells exhibit biochemical signatures of elevated PIP_2_ hydrolysis including higher baseline PM DAG-to PIP_2_ ratio and protein kinase C activity, and that PLC activity is essential for Pfn1-dependent changes in PM PIP_2_ content, supporting the idea of Pfn1’s regulation of PIP_2_ via impacting its hydrolysis in a cellular context. Fourth, we demonstrate for the first time that Pfn1 loss can have a major influence on cellular lipids that extend beyond PPIs.

Between the two cell lines we studied herein, Pfn1-dependent changes in PM PIP_2_ content were more prominent in MDA-231 cells (∼45%) relative to B16F1 (∼20%) cells. There could be at least two possible explanations for this observation. First, our MDA-231 studies were performed in acute KO setting, and this contrasts the experimental setting of B16F1 cells with prolonged Pfn1 loss where one cannot rule out the possibility of compensatory mechanisms. Second, we have confirmed that B16F1 cells have appreciably higher expression of Pfn2 (a minor isoform of Pfn) than MDA-231 cells (data not shown). Therefore, it is possible that increased presence of Pfn2 may partly offset the phenotypic changes associated with loss of Pfn1 in B16F1 cells.

Our conclusion that actin- rather than PPI-binding of Pfn1 is somehow responsible for Pfn1-dependent modulation of PIP_2_ is derived from functional rescue studies, where re-expression of all Pfn1 constructs except the actin-binding deficient mutant (H119E-Pfn1) rescued the PIP_2_ phenotype of Pfn1 KO cells. This was further supported by our LatB studies which revealed that F-actin loss (a natural consequence of loss of Pfn1) leads to rapid PIP_2_ loss at the PM in a temporally synchronized manner. Consistent with these results, we further showed that blebbistatin treatment resulted in accumulation of F-actin patches in cells and PIP_2_ enrichment in peripheral F-actin-rich regions. Together these findings demonstrate that PM PIP_2_ is significantly impacted by perturbation of actin cytoskeleton. However, given prior demonstration of Pfn1’s ability to protect PIP_2_ from PLC-mediated hydrolysis in model membrane settings (Goldschmidt-Clermont et al., 1991; Goldschmidt-Clermont et al., 1990), this begs a question as to whether we can completely disregard any contribution of Pfn1’s ability to modulate PIP_2_ by direct protein-lipid interaction in cells. First. it is noteworthy that in our previous study, we failed to see Pfn1’s recruitment to the PM even when we artificially increased PIP_2_ synthesis at the PM by chemical genetics strategy (Ricci et al., 2024). Since Pfn1 only displays transient, low-affinity interactions with PIP_2_-rich model membranes (Senju et al., 2017), we could not completely rule out technical limitations in capturing low abundant/affinity interactions between Pfn1:PIP_2_ at the PM in cells. Second, Pfn1 binds to actin and PPIs in a mutually exclusive manner. At least in a model membrane setting, Pfn1 can induce PIP_2_ clustering causing destabilization and deformation of the membrane (Krishnan et al., 2009). Therefore, one can logically argue that in a KO-rescue setting, H119E-Pfn1 has more frequent interactions with membrane PIP_2_ than either the WT or other mutant forms of Pfn1. As a result, local membrane deformation may render PIP_2_ more hydrolysis-prone partly contributing to reduced PM PIP_2_ content of the H119E-Pfn1 expressers. However, in our previous knockdown-rescue studies, we did not see any evidence for increased membrane content of H119E-Pfn1 mutant in MDA-231 cells (Bae et al., 2010). Therefore, the substantially diminished PM PIP_2_ content (in the range of 25-50% depending on the cell type) of Pfn1-deficient cells is very unlikely to be explained by the loss of low abundant/affinity Pfn1:PIP_2_ interaction, prompting us to speculate that the contribution of Pfn1’s ability to modulate PIP_2_ by direct protein-lipid interaction in cells, if at all, might be insignificant.

Although our whole-cell lipidomic studies revealed trends of elevated DAG and PA in Pfn1-deficient cells, because of greatest abundance of DAG in ER/Golgi compartments and metabolic fluxing of DAG to other lipids (TG, PA), those findings failed to provide granularity on whether Pfn1’s absence specifically alters PM PIP_2_ hydrolysis. However, our proposed tenet of Pfn1’s regulation of PIP_2_ via modulation of PIP_2_ hydrolysis is supported by multiple orthogonal lines of evidence. First, our lipid biosensor studies revealed greater PM DAG-to-PIP_2_ ratio (a measure of the intrinsic PIP_2_ hydrolysis efficiency) in Pfn1-deficient cells. Second, GSEA of transcriptomic data predicted enrichment of cytosolic IP_3_/IP_4_ synthesis pathway in Pfn1-deficient cells. Third, cells lacking Pfn1 exhibited increased baseline PKC activity. Fourth, PLC inhibition resulted in reversal of Pfn1-dependent changes in PIP_2_. Two previous have demonstrated that actin filament disruption increases PM mobility of PIP_2_ (Andrade et al., 2015; Cho et al., 2005). There is also evidence for actin depolymerization-induced uncaging of PLC from the cortical actin network (Huang and Crain, 2009). Therefore, it is possible that Pfn1 deficiency leads to loosening of cortical actin network leading to increased PM diffusion of PIP_2_ and/or uncaging of PLC, and this results in more frequent PLC-PIP_2_ interaction and higher baseline PIP_2_ hydrolysis, as schematically represented in **Figure 7**.

**Figure 7:**
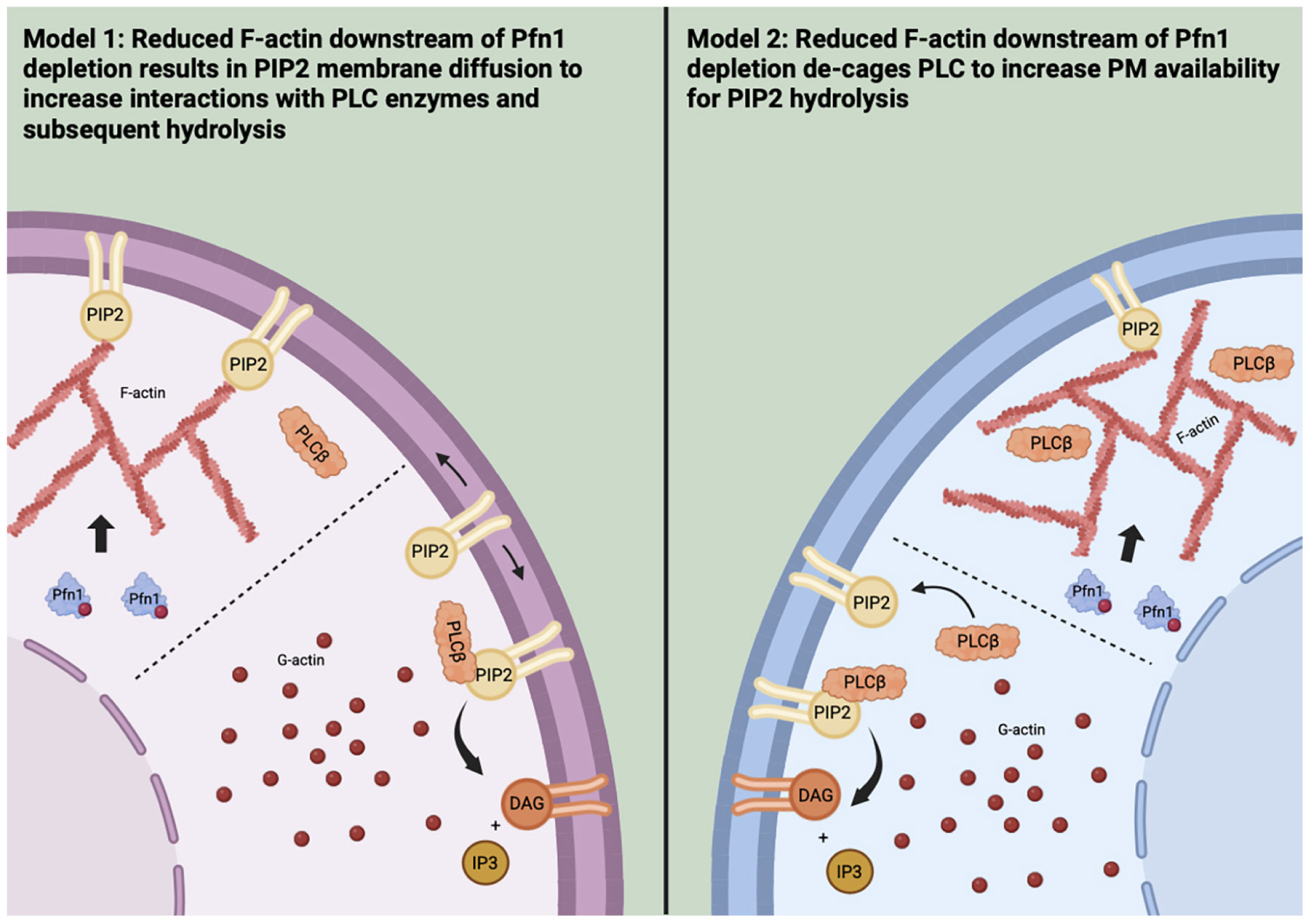
Proposed schematic models of how Pfn1 loss could potentially accelerate PIP_2_ hydrolysis in an actin-dependent manner (see main text for detailed explanation).

Although Pfn1 depletion has no discernible impact on PM PI4P (the precursor PPI for PIP_2_ synthesis) content (this is also consistent with of comparable PI content between control and Pfn1 KO cells), a limitation of the present study is that we have not explored the possibility of Pfn1-dependent alterations in localization and/or activity of cellular machinery (i.e. PIP5K) for PIP_2_ biosynthesis. For example, a previous study reported that Pfn1 depletion in chondrocytes inhibits Rho GTPase activation (Bottcher et al., 2009). Since the enzymatic activity of PIP5K is stimulated by activated Rho (Weernink et al., 2004), it is not outside of the realm of possibility that Pfn1 loss may still have a negative impact on the actual enzymatic activity of PIP5K via inhibition of Rho. Alternatively, loss of Pfn1 may influence the PM localization of PIP5K (critical for PIP_2_ synthesis) as we had previously shown for SHIP2 (Ricci et al., 2024). Even if some of these possibilities are true, this is still unlikely to be the dominant mechanism given that inhibition of PIP_2_ hydrolysis pathway rescues the PIP_2_ phenotype of Pfn1-deficient cells.

Finally, our proof-of-concept lipidomic profiling shows that Pfn1 modulates a broader range of lipid species that extend beyond PPIs in cells revealing a previously unrecognized role of Pfn1 in lipid homeostasis. Of particular importance, we found evidence for prominent decrease in PC, cholesterol and sphingomyelin, and increase in TG in cells when Pfn1 expression is suppressed. PC is the most abundant phospholipid making up 40-50% of the phospholipid content in cells and plays a major role in determining cell membrane structure (van der Veen et al., 2017). Cholesterol and sphingolipids are major molecular constituents of lipid rafts, the microdomains in the cell membrane that serve as platforms for protein localization and signaling hotspots (Lingwood and Simons, 2010). TG plays critical functions as energy reservoirs and its hydrolysis via lipolysis into free fatty acids that are subsequently oxidized in mitochondria to generate ATP (Kwiterovich, 2000). Therefore, we think our lipidomic findings open completely new directions for future studies investigating whether and how Pfn1 regulates various biological processes via lipid control. Those findings could shed novel mechanistic insights linking Pfn1 dysregulation to disease progression.

## MATERIALS AND METHODS

### Cell Culture, plasmids, and siRNA transfection

HEK-293A cells (Thermo Fisher Scientific R705-07) were cultured in complete media composed of low glucose Dulbecco’s modified Eagle’s medium (DMEM) (Thermo Fisher Scientific 10567022), 10% heat-inactivated fetal bovine serum (Thermo Fisher Scientific 10438-034), 100 μg/ml penicillin, 100 μg/ml streptomycin (Thermo Fisher Scientific #15140122), and 0.1% chemically defined lipid supplement (Thermo Fisher Scientific #11905031). GFP/Luciferase expressing sublines of MDA-MB-231 (MDA-231 – source: ATCC) breast cancer cells (as described previously (Gau et al.)) were cultured in in DMEM supplemented with 10% (v/v) fetal bovine serum, 100 μg/ml penicillin, 100 μg/ml streptomycin, and geneticin (100 μg/ml). To generate doxycycline-inducible Pfn1 knockout (KO) MDA-231 cell line, we utilized the Edit-R Inducible Lentiviral CRISPR/Cas9 system following manufacturer’s protocol (Pfn1 sgRNA: VSGH10142-246635145, non-targeting sgRNA: VSGC10215, Cas9 nuclease: VCAS11227, Horizon). MOI of 0.3 was utilized for sequential transduction of Cas9- and gRNA-encoding lentivirus for stable cell generation. Cas9 positive clones were selected and maintained with blasticidin (100 μg/mL). Pfn1 or non-targeting KO clones were selected and maintained puromycin (2.5 μg/mL). Doxycycline was prepared at 1mg/mL and administered to cells at a final concentration of 1μg/mL to trigger Pfn1 KO and experimental analyses were performed 6 days after initial doxycycline induction. Generation and culture of stable Pfn1 KO B16F1 melanoma cells have been described elsewhere (Tang et al., 2025). All cell lines were routinely screened for confirmation of mycoplasma-free status by PCR. Lipofectamine 3000 and RNAi Max were used for plasmid and siRNA transfection of cells, respectively, as per manufacturer’s instructions. Various myc-tagged Pfn1 constructs were cloned into an IRES-GFP backbone vector. For transient silencing studies, siRNAs targeting Pfn1 (as previously described (Joy et al., 2017)) and PLCβ3 (Thermo Fisher #AM16708) were transfected in cells and downstream analyses were performed at least 72 hrs after transfection. For intracellular Ca^2+^ measurement, live imaging of cells transfected with GCaMP, a genetically encoded Ca^2+^ sensor (source: Addgene), was performed.

### PPI immunostaining and quantification

For PIP_2_ immunostaining, cells were seeded on glass substrates coated with either rat tail collagen or murine laminin 24 hours prior to staining. Cells were rapidly fixed in 4% formaldehyde and 0.2% glutaraldehyde fixation buffer for 15 min at room temperature (20–24°C) before rinsing three times with PBS containing 50 mM NH_4_Cl. Cells were then chilled on ice for at least 10 minutes before proceeding. All subsequent steps were performed on ice, with all solutions pre-chilled. Cells were blocked and permeabilized for 45 min with a solution of buffer A containing PBS with 5% (v/v) normal goat serum (NGS), 50 mM NH_4_Cl and 0.5% saponin, before incubation with a 1:100 dilution of mouse monoclonal anti-PIP_2_ antibody (Echelon Biosciences; Catalog #Z-PO45) in buffer B containing PBS with 5% NGS and 0.1% saponin. Note that the selectivity of this PIP_2_ antibody was previously established in immunostaining studies where the immunofluorescence signal was lost by either antibody-competing PIP_2_ or neomycin (binds to PIP_2_ with high affinity) or the pleckstrin-homology domain of PIP_2_-binding protein PLCδ (Hammond et al., 2006; Hammond et al., 2009). At the end of primary antibody incubation, cells were washed three washes in buffer B before incubation with a secondary antibody in buffer B for 45 minutes followed by four washes in buffer B. Cells were then post-fixed in 2% formaldehyde in PBS for 10 min on ice, before warming to room temperature for an additional 5 min. Formaldehyde was removed by three rinses in PBS containing 50 mM NH_4_Cl, followed by one rinse in PBS. Fluorescence images were acquired on an IX83 Olympus (with Cicero confocal) microscope using either a 20x or 40x objective. For PI4P immunostaining, a mouse monoclonal anti-PI(4)P (Echelon Biosciences; Catalog #Z-P004; 1:100 dilution) antibody was used with identical procedures.

To analyze plasma membrane (PM) fluorescence intensity, fluorescence images were saved as .tif files and processed using CellProfiler image analysis software. Background subtraction was performed by generating an illumination correction function, smoothing it with a Gaussian blur, and subtracting it from the original image. Regions of interest (cell boundaries) were identified by an Otsu threshold algorithm for quantifying the integrated fluorescence intensities at the edge for each cell. For functional rescue experiments, this pipeline was supplemented with additional steps to identify vector expressing cells by saving multi-channel fluorescence images (GFP and RFP), manually segmenting GFP-positive cells, and then employing a distance-based propagation method to define ROIs corresponding to cells co-expressing both fluorophores and cell boundaries prior to integrated edge intensity measurement.

### Lipid biosensor studies

HEK-293, co-transfected with Tubby_c-_EGFP (a biosensor for PIP_2_), mCherry-C1ab-PKD1 (a biosensor for DAG) and iRFP-Lyn11 (a PM biomarker), were imaged with a confocal microscope for quantification of PM to cytosolic fluorescence intensities for various reporters.

### F-actin staining and quantification

F-actin staining was performed by fixing cells in 3.7% formaldehyde for 15 minutes at 37 C. After 3 washes with DPBS, samples were permeabilized with 0.5% triton X in PBS at room temperature for 5 minutes. Following another 3 washes with DPBS, samples were incubated with 1:400 FITC-Phalloidin Stain (Sigma-Aldrich: P5282) in PBS for 1 hour at room temperature in the dark, after which the staining solution was aspirated, and samples were washed 3x with PBS. The samples were imaged using either 20x or 60x objectives. For live imaging of F-actin, cells were labeled with SPY55FastActX (Cytoskeleton Inc) as per the manufacturer’s instruction.

### Pharmacological studies

In experiments involving pan-PLC inhibitor U73122 and its inactive control U73343, the dry compounds were diluted in DMSO to a stock concentration of 10mM and added to cells at a final concentration of 2uM after 24 hours of serum stimulation. In experiments using Lysophosphatidic Acid (LPA), the agonist was diluted to a stock concentration of 10mM and administered to cells after 24 hours of serum starvation at a final concentration of 10μM for 5 minutes prior to or during experimental analysis. For studies using latrunculin B (LatB), the compound was diluted to a stock concentration of 1mM and administered to cells at a working concentration of 2 μM for 5 minutes prior to analysis. For studies involving inhibition of myosin, blebbistatin was diluted to stock concentrations of 10mM and added to the cultures at a final concentration of 50μM for 30 minutes prior to end-point analysis.

### Lipidomic Analysis

Quantitative lipidomic analysis was performed by Metaware Bio (Durham, NC, USA). Cells were grown to 80% confluency at which point they were trypsinized, washed 2x with PBS, and aliquoted into pellets of >1,000,000 cells per sample and immediately flash-frozen in liquid nitrogen and stored at −80°C until shipment. Upon receipt, Metaware Bio conducted comprehensive lipidomic profiling using liquid chromatography–mass spectrometry (LC-MS/MS). A detailed protocol of their quantitative lipidomic panel can be found at (https://www.metwarebio.com/lipidomics/).

### Immunoblot

Total cell lysate was collected using a modified RIPA buffer (25 mM Tris–HCl, pH 7.5, 150 mM NaCl, 1% (v/v) Nonidet P-40, 0.2% SDS, 5% (v/v) glycerol, 1 mM EDTA) supplemented with protease/phosphatase inhibitor cocktail. Total cell lysate was run on an SDS-PAGE and immunoblotted using various antibodies: Pfn1 monoclonal antibody (Abcam, #EPR6304, 1:750), PLCβ3 polyclonal antibody (Invitrogen, #PA5-78109; 1:1000), GAPDH monoclonal antibody (Invitrogen, #MAS-15738; 1:2000) and Phospho(ser)PKC substrate polyclonal antibody (Cell signaling technology, #2261; 1:1000).

### Bulk transcriptome sequencing (RNA-seq)

For transcriptome sequencing, total RNA was extracted from MDA-231 cells (control vs Pfn1 KO) using a commercial kit (Qiagen). The mRNA was isolated and reverse-transcribed to cDNA and amplified using Illumina® Stranded mRNA Prep from Illumina, Inc (San Diego, CA). The library preparation processes such as adenylation, ligation, and amplification were performed following the manual provided by the manufacturer. The quantity and quality of the libraries were assessed through Bioanalyzer 2100 and Qubit instruments. The procedure of 150 cycle paired-end sequencing in AVITI sequencer followed the manufacturer’s manual (Love et al., 2014). Approximately 25 million sequencing reads were generated per sample, and these reads were trimmed using Trimmomatic to filter out low-quality reads and adapter sequences (Bolger et al., 2014). The surviving reads were then aligned to the reference genome, and the gene-level counts were quantified using STAR with the quantMode GeneCounts option. Differential expression analysis was performed on the gene-by-sample count matrix using the DESeq2 package in R/Bioconductor. Differentially expressed genes (DEGs) were defined by an FDR equal to 5% and an absolute fold change > 1.5. These DEGs were subsequently used for gene set enrichment analyses (GSEA).

### Statistics

All statistical tests were performed using GraphPad Prism 9 software (https://www.graphpad.com). ANOVA (for more than two groups) and Student’s T test (for two groups) were performed for comparing means between various experimental groups. A *p*-value less than 0.05 was statistically significant. All error bars in the figures represent standard deviations of data unless explicitly stated otherwise.

## AUTHOR CONTRIBUTIONS

AO performed experiments, analyzed data, and wrote the manuscript; MC performed experiments and analyzed data, IE performed experiments, YT and DG contributed to generation of technical reagents, JL analyzed data, SL, JL, KR, and GH oversaw experiments and data analyses, PR wrote/edited the manuscript, acquired funding, and was responsible for overall supervision of the project.

## ACKNOWLEDGEMENTS

Research in the Roy lab was supported by funding from the National Institute of Health (NIH) grants R01CA248873, R01CA271095, R21EY-032632 and Department of Defense grant HT425-24-2-0556.

## DATA AVAILABILITY

All relevant data and details of resources can be found within the article and its supplementary information.

## REFERENCES

Akiyama, C., Shinozaki-Narikawa, N., Kitazawa, T., Hamakubo, T., Kodama, T. and Shibasaki, Y. (2005). Phosphatidylinositol-4-phosphate 5-kinase gamma is associated with cell-cell junction in A431 epithelial cells. Cell Biol Int 29, 514–20.

Andrade, D. M., Clausen, M. P., Keller, J., Mueller, V., Wu, C., Bear, J. E., Hell, S. W., Lagerholm, B. C. and Eggeling, C. (2015). Cortical actin networks induce spatio-temporal confinement of phospholipids in the plasma membrane--a minimally invasive investigation by STED-FCS. Sci Rep 5, 11454.

Bae, Y. H., Ding, Z., Das, T., Wells, A., Gertler, F. and Roy, P. (2010). Profilin1 regulates PI(3,4)P2 and lamellipodin accumulation at the leading edge thus influencing motility of MDA-MB-231 cells. Proc Natl Acad Sci U S A 107, 21547–52.

Bolger, A. M., Lohse, M. and Usadel, B. (2014). Trimmomatic: a flexible trimmer for Illumina sequence data. Bioinformatics 30, 2114–20.

Bottcher, R. T., Wiesner, S., Braun, A., Wimmer, R., Berna, A., Elad, N., Medalia, O., Pfeifer, A., Aszodi, A., Costell, M. et al. (2009). Profilin 1 is required for abscission during late cytokinesis of chondrocytes. EMBO J.

Braun, L., Schoen, I. and Vogel, V. (2021). PIP(2)-induced membrane binding of the vinculin tail competes with its other binding partners. Biophys J 120, 4608–4622.

Chinthalapudi, K., Rangarajan, E. S., Patil, D. N., George, E. M., Brown, D. T. and Izard, T. (2014). Lipid binding promotes oligomerization and focal adhesion activity of vinculin. J Cell Biol 207, 643–56.

Cho, H., Kim, Y. A., Yoon, J. Y., Lee, D., Kim, J. H., Lee, S. H. and Ho, W. K. (2005). Low mobility of phosphatidylinositol 4,5-bisphosphate underlies receptor specificity of Gq-mediated ion channel regulation in atrial myocytes. Proc Natl Acad Sci U S A 102, 15241–6.

Czech, M. P. (2000). PIP2 and PIP3: complex roles at the cell surface. Cell 100, 603–6.

Davey, R. J. and Moens, P. D. (2020). Profilin: many facets of a small protein. Biophys Rev 12, 827–849.

Dickson, E. J. and Hille, B. (2019). Understanding phosphoinositides: rare, dynamic, and essential membrane phospholipids. Biochem J 476, 1–23.

Ding, Z., Bae, Y. H. and Roy, P. (2012). Molecular insights on context-specific role of profilin-1 in cell migration. Cell Adh Migr 6, 442–9.

Gau, D., Chawla, P., Eder, I. and Roy, P. Myocardin-related transcription factor’s interaction with serum-response factor is critical for outgrowth initiation, progression, and metastatic colonization of breast cancer cells. FASEB BioAdvances n/a.

Goldschmidt-Clermont, P. J., Kim, J. W., Machesky, L. M., Rhee, S. G. and Pollard, T. D. (1991). Regulation of phospholipase C-gamma 1 by profilin and tyrosine phosphorylation. Science 251, 1231–3.

Goldschmidt-Clermont, P. J., Machesky, L. M., Baldassare, J. J. and Pollard, T. D. (1990). The actin-binding protein profilin binds to PIP2 and inhibits its hydrolysis by phospholipase C. Science 247, 1575–8.

Hammond, G. R., Dove, S. K., Nicol, A., Pinxteren, J. A., Zicha, D. and Schiavo, G. (2006). Elimination of plasma membrane phosphatidylinositol (4,5)-bisphosphate is required for exocytosis from mast cells. J Cell Sci 119, 2084–94.

Hammond, G. R., Schiavo, G. and Irvine, R. F. (2009). Immunocytochemical techniques reveal multiple, distinct cellular pools of PtdIns4P and PtdIns(4,5)P(2). Biochem J 422, 23–35.

Harraz, O. F., Hill-Eubanks, D. and Nelson, M. T. (2020). PIP(2): A critical regulator of vascular ion channels hiding in plain sight. Proc Natl Acad Sci U S A 117, 20378–20389.

Higgs, H. N. and Pollard, T. D. (2000). Activation by Cdc42 and PIP(2) of Wiskott-Aldrich syndrome protein (WASp) stimulates actin nucleation by Arp2/3 complex. J Cell Biol 150, 1311–20.

Huang, C. H. and Crain, R. C. (2009). Phosphoinositide-specific phospholipase C in oat roots: association with the actin cytoskeleton. Planta 230, 925–33.

James, J., Fokin, A. I., Guschin, D. Y., Wang, H., Polesskaya, A., Rubtsova, S. N., Clainche, C. L., Silberzan, P., Gautreau, A. M. and Romero, S. (2025). Vinculin-Arp2/3 interaction inhibits branched actin assembly to control migration and proliferation. Life Sci Alliance 8.

Joy, M., Gau, D., Castellucci, N., Prywes, R. and Roy, P. (2017). The myocardin-related transcription factor MKL co-regulates the cellular levels of two profilin isoforms. J Biol Chem 292, 11777–11791.

Krishnan, K., Holub, O., Gratton, E., Clayton, A. H., Cody, S. and Moens, P. D. (2009). Profilin interaction with phosphatidylinositol (4,5)-bisphosphate destabilizes the membrane of giant unilamellar vesicles. Biophys J 96, 5112–21.

Kwiterovich, P. O., Jr. (2000). The metabolic pathways of high-density lipoprotein, low-density lipoprotein, and triglycerides: a current review. Am J Cardiol 86, 5l–10l.

Lambrechts, A., Verschelde, J. L., Jonckheere, V., Goethals, M., Vandekerckhove, J. and Ampe, C. (1997). The mammalian profilin isoforms display complementary affinities for PIP2 and proline-rich sequences. EMBO J 16, 484–94.

Lassing, I. and Lindberg, U. (1985). Specific interaction between phosphatidylinositol 4,5-bisphosphate and profilactin. Nature 314, 472–4.

Lee, S., Park, J. B., Kim, J. H., Kim, Y., Kim, J. H., Shin, K. J., Lee, J. S., Ha, S. H., Suh, P. G. and Ryu, S. H. (2001). Actin directly interacts with phospholipase D, inhibiting its activity. J Biol Chem 276, 28252–60.

Lingwood, D. and Simons, K. (2010). Lipid rafts as a membrane-organizing principle. Science 327, 46–50.

Love, M. I., Huber, W. and Anders, S. (2014). Moderated estimation of fold change and dispersion for RNA-seq data with DESeq2. Genome Biol 15, 550.

Lu, P. J., Shieh, W. R., Rhee, S. G., Yin, H. L. and Chen, C. S. (1996). Lipid products of phosphoinositide 3-kinase bind human profilin with high affinity. Biochemistry 35, 14027–34.

Mandal, K. (2020). Review of PIP2 in Cellular Signaling, Functions and Diseases. Int J Mol Sci 21.

Mondal, S., Pal, B. and Sankaranarayanan, R. (2024). Diacylglycerol metabolism and homeostasis in fungal physiology. FEMS Yeast Res 24.

Ricci, M. M. C., Orenberg, A., Ohayon, L., Gau, D., Wills, R. C., Bae, Y., Das, T., Koes, D., Hammond, G. R. V. and Roy, P. (2024). Actin-binding protein profilin1 is an important determinant of cellular phosphoinositide control. J Biol Chem 300, 105583.

Senju, Y., Kalimeri, M., Koskela, E. V., Somerharju, P., Zhao, H., Vattulainen, I. and Lappalainen, P. (2017). Mechanistic principles underlying regulation of the actin cytoskeleton by phosphoinositides. Proc Natl Acad Sci U S A 114, E8977–e8986.

Shutova, M., Yang, C., Vasiliev, J. M. and Svitkina, T. (2012). Functions of nonmuscle myosin II in assembly of the cellular contractile system. PLoS One 7, e40814.

Sohn, R. H., Chen, J., Koblan, K. S., Bray, P. F. and Goldschmidt-Clermont, P. J. (1995). Localization of a binding site for phosphatidylinositol 4,5-bisphosphate on human profilin. J Biol Chem 270, 21114–20.

Tang, Y., Schaks, M., Mietkowska, M., Scholz, J., Benavente-Naranjo, R., Körber, S., Li, Z., Lambert, C., Stradal, T. E. B., Karlsson, R. et al. (2025). Profilin promotes lamellipodium protrusion by tuning the antagonistic activities of CP and VASP. BioRxiv, 2025.03.26.645585.

Thapa, N. and Anderson, R. A. (2012). PIP2 signaling, an integrator of cell polarity and vesicle trafficking in directionally migrating cells. Cell Adh Migr 6, 409–12.

van der Veen, J. N., Kennelly, J. P., Wan, S., Vance, J. E., Vance, D. E. and Jacobs, R. L. (2017). The critical role of phosphatidylcholine and phosphatidylethanolamine metabolism in health and disease. Biochim Biophys Acta Biomembr 1859, 1558–1572.

Van Veldhoven, P. P. and Bell, R. M. (1988). Effect of harvesting methods, growth conditions and growth phase on diacylglycerol levels in cultured human adherent cells. Biochim Biophys Acta 959, 185–96.

Weernink, P. A., Meletiadis, K., Hommeltenberg, S., Hinz, M., Ishihara, H., Schmidt, M. and Jakobs, K. H. (2004). Activation of type I phosphatidylinositol 4-phosphate 5-kinase isoforms by the Rho GTPases, RhoA, Rac1, and Cdc42. J Biol Chem 279, 7840–9.

Ye, X., McLean, M. A. and Sligar, S. G. (2016). Phosphatidylinositol 4,5-Bisphosphate Modulates the Affinity of Talin-1 for Phospholipid Bilayers and Activates Its Autoinhibited Form. Biochemistry 55, 5038–48.

Zhao, H., Hakala, M. and Lappalainen, P. (2010). ADF/cofilin binds phosphoinositides in a multivalent manner to act as a PIP(2)-density sensor. Biophys J 98, 2327–36.

