## Supplementary Info for "Molecular insights into Profilin1-dependent regulation of cellular phosphatidylinositol-(4,5)-bisphosphate"

**Supplementary Information** (Orenberg et al.)

Orenberg et al. Fig S1

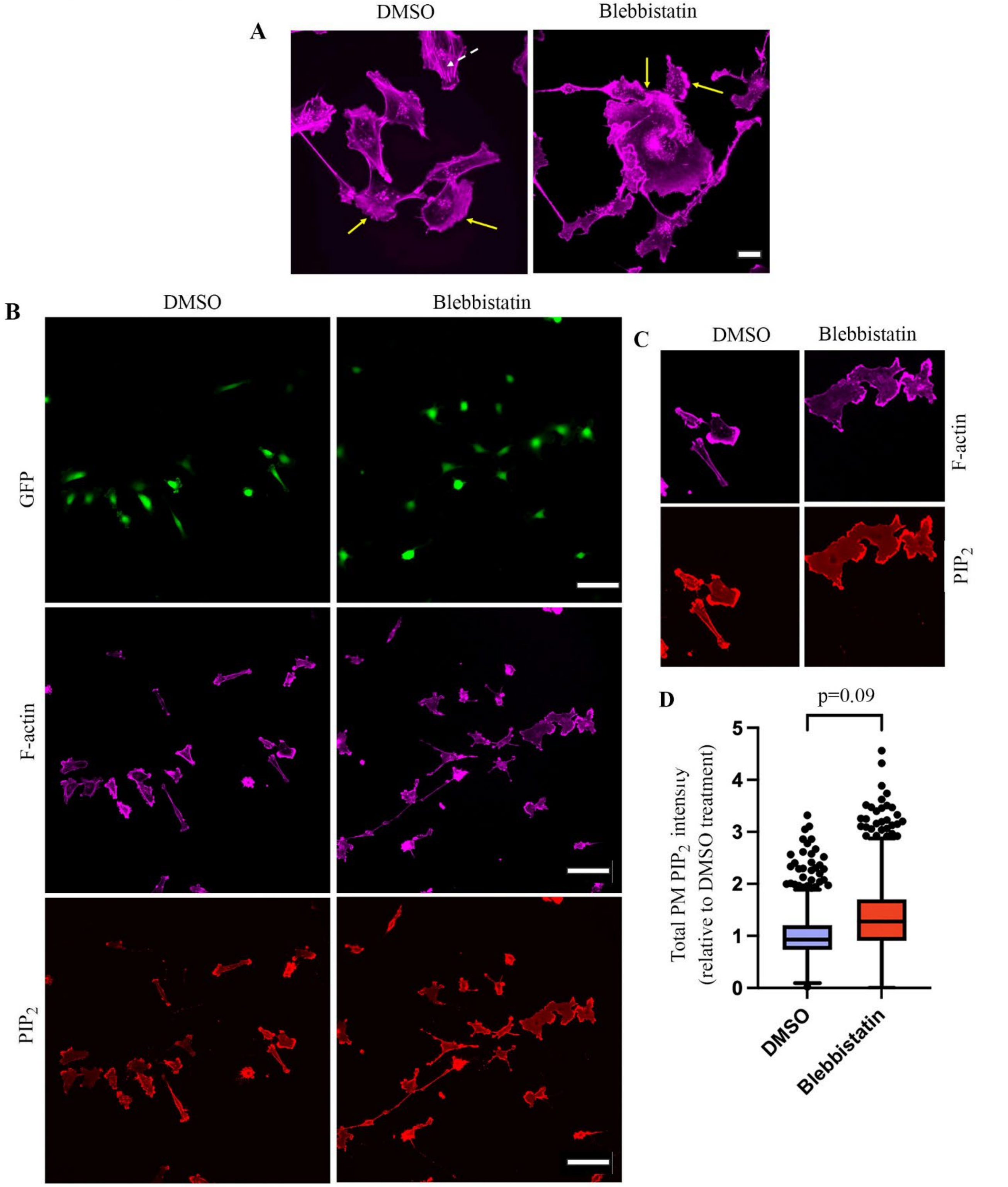


**Fig S1: Effect of blebbistatin treatment on F-actin- and $PIP_2$ in MDA-231 cells. A)** Representative phalloidin staining images (60X magnification) of GFP-expressing subline of MDA-231 cells subjected to 30 min treatment of either blebbistatin or DMSO (scale bar – 20 $\mu$m). Arrowheads show actin stress fibers in DMSO-treated cells that were largely eliminated upon blebbistatin treatment. Arrows indicate peripheral F-actin. **B-C)** Representative widefield fluorescence images (20X magnification; scale bar – 40 $\mu$m) of phalloidin- and $PIP_2$-stained GFP-expressing MDA-231 cells (panel B; magnified images of F-actin and $PIP_2$ staining of selected regions of interest are shown in panel C). **D)** A box and whisker plot summarizing the relative total cell edge $PIP_2$ staining intensity of the two treatment groups (data summarized from analyses of more than 700 individual cells per treatment group pooled from 3 experiments).

Orenberg et al. Fig S2

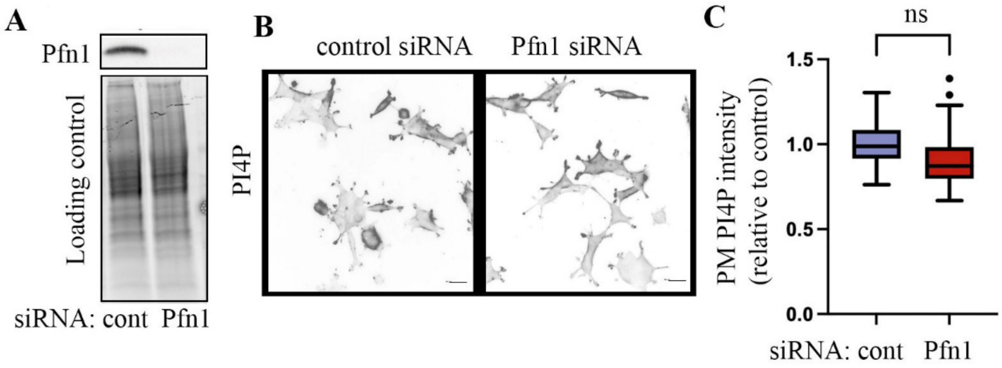


**Fig S2: Pfn1 loss does not impact PI4P content in HEK-293 cells. A)** Pfn1 immunoblot (stain-free gel serves as the loading control) of HEK-293 cells transfected with either control or Pfn1-specific siRNAs show efficient knockdown of Pfn1 expression. **B-C)** Representative images of PI4P immunostaining (*panel B;* scale – 50 $\mu$m) and quantification (*panel C*) of control vs Pfn1 knockdown cultures (data summarized from 300-400 cells/group pooled from 3 experiments; ns – not significant).

Orenberg et al. Fig S3

A

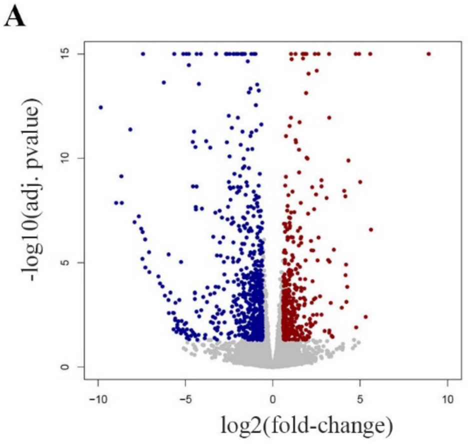


B

| DOWNREGULATED | | | UPREGULATED | |
|---|---|---|---|---|
| Gene | log2FoldChange | | Gene | log2foldchange |
| FOXP2 | -9.85 | | LOC10192 | 12.68 |
| NGFR | -8.97 | | CMKLR1 | 8.92 |
| CDH11 | -8.67 | | CHRNE | 5.62 |
| FAM49A | -8.65 | | AKR1C1 | 5.58 |
| CACNA2D | -8.16 | | KCNJ4 | 5.32 |
| GABRA3 | -7.92 | | ZBTB32 | 5.00 |
| VCAN | -7.67 | | MAGEA11 | 4.94 |
| TRPM6 | -7.54 | | LINC0122 | 4.77 |
| ZNF804A | -7.47 | | AKR1C2 | 4.75 |
| AMOT | -7.45 | | C17orf107 | 4.33 |
| CDK14 | -7.43 | | RSAD2 | 4.26 |
| FREM2 | -7.31 | | KRT17 | 4.21 |
| BCHE | -7.30 | | SORCS3 | 4.19 |
| SLITRK1 | -7.09 | | CCL5 | 4.16 |
| TRIML2 | -7.07 | | ELMO1 | 4.15 |
| TCEAL7 | -6.55 | | NAP1L3 | 4.09 |
| MAGEA10 | -6.46 | | IFNL1 | 3.94 |
| COCH | -6.32 | | CES1P2 | 3.49 |
| GYPC | -6.24 | | LINC0064 | 3.42 |
| LDOC1 | -6.18 | | CH25H | 3.37 |
| C3orf70 | -5.97 | | PLXNA4 | 3.31 |
| PAGE5 | -5.97 | | ITGAX | 3.24 |
| RASEF | -5.95 | | FA2H | 3.24 |
| FGF13 | -5.86 | | NCF2 | 3.23 |
| MARK1 | -5.71 | | LOC10192 | 3.22 |
| SHROOM4 | -5.69 | | AKR1C3 | 3.22 |
| LINC0026 | -5.67 | | ZBTB7C | 3.17 |
| GPC6 | -5.64 | | C7orf71 | 3.08 |
| PTPRO | -5.59 | | RTP4 | 2.96 |
| LINC0132 | -5.59 | | MINK1 | 2.79 |
| KANK1 | -5.58 | | MAFIP | 2.79 |
| SLCO4C1 | -5.52 | | LINC0070 | 2.78 |
| LRAT | -5.48 | | TONSL-AS | 2.70 |
| LYPD6B | -5.47 | | CST1 | 2.62 |
| ELMOD1 | -5.43 | | BIK | 2.61 |
| AMPH | -5.41 | | PI16 | 2.56 |
| FAM184A | -5.41 | | SLC22A18 | 2.52 |
| ENPP5 | -5.36 | | NAA11 | 2.51 |
| PTGFR | -5.34 | | CEACAM1 | 2.47 |
| CRISPLD1 | -5.26 | | GCKR | 2.44 |
| PTGFRN | -5.26 | | PRSS36 | 2.44 |
| MIR137HG | -5.22 | | RAB40C | 2.41 |
| CSRNP3 | -5.17 | | UNC13A | 2.36 |
| NDNF | -5.15 | | OASL | 2.35 |
| MAGEA6 | -5.13 | | GPR132 | 2.31 |
| MIR99AHG | -5.06 | | GIMAP6 | 2.30 |
| RERG | -5.05 | | NAALAD2 | 2.28 |
| ANGPT1 | -4.97 | | C1QTNF1- | 2.27 |
| ZNF300 | -4.97 | | TRIM29 | 2.25 |
| ARHGDIB | -4.95 | | NEURL3 | 2.23 |

C

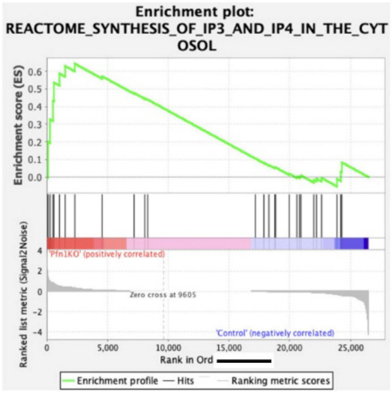


D

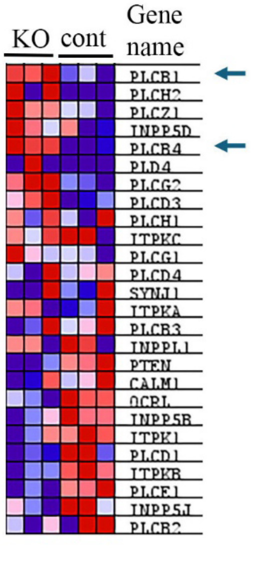


**Fig S3: Summary of transcriptomic findings of control vs Pfn1 KO MDA-231 cells: A)** Volcano plot representing differentially expressed genes between control and Pfn1 KO MDA-231 cells (upon Pfn1 KO in MDA-231 cells. **B)** List of top 100 (50 up- and 50-downregulatde) differentially expressed genes in Pfn1 KO relative to control culture. **C-D)** GSEA analyses of transcriptome data (*panel C*) predicts enrichment (normalized enrichment score -1.4; nominal p-value – 0.0) of cytosolic $IP_3$/$IP_4$ synthesis-related genes in Pfn1 KO MDA-231 cells; *panel D* shows the associated heat-plot of gene expression (the top 7 listed genes contribute to the enrichment; arrows indicate the PLC isoform genes that are transcriptionally upregulated (~4,0 fold-increase) in a statistically significant manner in Pfn1 KO cells vs their control counterparts.
